# Modelling Human Post-Implantation Development via Extra-Embryonic Niche Engineering

**DOI:** 10.1101/2023.06.15.545118

**Authors:** Joshua Hislop, Amir Alavi, Qi Song, Rayna Schoenberger, Kamyar Keshavarz F., Ryan LeGraw, Jeremy Velazquez, Tahere Mokhtari, Mohammad Nasser Taheri, Matthew Rytel, Susana M Chuva de Sousa Lopes, Simon Watkins, Donna Stolz, Samira Kiani, Berna Sozen, Ziv Bar-Joseph, Mo R. Ebrahimkhani

## Abstract

Implantation of the human embryo commences a critical developmental stage that comprises profound morphogenetic alteration of embryonic and extra-embryonic tissues, axis formation, and gastrulation events. Our mechanistic knowledge of this window of human life remains limited due to restricted access to *in vivo* samples for both technical and ethical reasons. Additionally, human stem cell models of early post-implantation development with both embryonic and extra-embryonic tissue morphogenesis are lacking. Here, we present iDiscoid, produced from human induced pluripotent stem cells via an engineered a synthetic gene circuit. iDiscoids exhibit reciprocal co-development of human embryonic tissue and engineered extra-embryonic niche in a model of human post-implantation. They exhibit unanticipated self-organization and tissue boundary formation that recapitulates yolk sac-like tissue specification with extra-embryonic mesoderm and hematopoietic characteristics, the formation of bilaminar disc-like embryonic morphology, the development of an amniotic-like cavity, and acquisition of an anterior-like hypoblast pole and posterior-like axis. iDiscoids offer an easy-to-use, high-throughput, reproducible, and scalable platform to probe multifaceted aspects of human early post-implantation development. Thus, they have the potential to provide a tractable human model for drug testing, developmental toxicology, and disease modeling.

## Main Text

Decoding mechanisms of human embryogenesis has tremendous biomedical impact, from treating congenital diseases and infertility, to engineering functional human organs. Immediately after implantation, the embryo and co-developing extra-embryonic tissues are profoundly remodeled and initiate morphological changes central to the success of pregnancy, including formation of the amniotic cavity and emergence of yolk sac hematopoiesis^1, 2^. However, hidden within the uterine tissues, the key steps of early post-implantation development in humans largely remain beyond our reach, owing to limited access to this stage for both technical and ethical reasons^3, 4^. The early post-implantation stage human embryo, with its flat bilaminar disc structure, is morphologically different from its mouse counterpart. As such it may demand new engineering approaches to be efficiently modeled. *In vitro* models of embryogenesis based on both mouse and human-derived cell types have emerged as instrumental platforms to probe human-specific developmental mechanisms^5–20^. However, they often suffer limitations, such as low efficiency and high technical complexity. Traditionally these studies use cocktails of growth factors at supra-physiological levels and encounter challenges finding common media that support diverse cellular fates. Additionally, human post-implantation embryo models with co-developing embryonic and extra-embryonic layers are still missing, and a full integrated human embryo-like model can encounter ethical challenges, which limits our ability to study integrated morphogenetic events of human embryo after implantation^3, 21^.

Here, we present a human stem cell-derived embryo model, dubbed “***iDiscoid***”, that overcomes several of these limitations. The iDiscoid platform leverages an approach that genetically engineers a population of human induced pluripotent stem cells (hiPSCs) with inducible circuits. Upon activation of these genetic circuits, an extra-embryonic niche is formed that triggers self-organization of the adjacent wild-type hiPSC clusters. Using iDiscoids, we demonstrate development of human bilaminar disc-like structures, specification of anterior-like hypoblast cells, lumenogenesis, formation of amnion-like tissue, symmetry breaking leading to posterior-like axis specification, and yolk sac specification with hematopoietic characteristics. iDiscoids model human embryogenesis at an early stage of implantation when the embryo *in vivo* is at its most inaccessible state.

### Engineering co-development of embryonic and extra-embryonic endoderm tissues

We engineered an hiPSC cell line with an inducible gene circuit that expresses human GATA6, a key transcription factor for extra-embryonic endodermal fate (henceforth referred to as iGATA6) (**Fig. 1A, Supplementary Fig. 1A)**. These iGATA6 cells express different levels of GATA6 upon the addition of a small molecule, Doxycycline (Dox), distinguishable by expression of the reporter enhanced green fluorescent protein (EGFP) **(Supplementary Fig. 1B)**. When iGATA6 and wild-type (WT) hiPSCs were co-assembled in 3D, EGFP^+^ extra-embryonic endoderm cells segregated to the outer layer, similar to what occurs during implantation period *in vivo.* However, the overall organization of these aggregates did not recapitulate the structural format of bilaminar disc and yolk sac tissue. This approach yielded aggregates in which the entire EGFP^+^ extra-embryonic endoderm is in close contact with wild type epiblast-like cells, missing the part of the yolk sac tissue (parietal) that is formed away from the epiblast cells. Additionally, in these 3D aggregates, we found that WT cells could not consistently induce symmetry breaking to amnion-like and epiblast-like domains **(Supplementary Fig. 1C-D)**. Moreover, *in vitro* reattachment of both synthetic and true human blastocysts to a dish to mimic implantation has shown organizational instability that results in tissue organization that cannot reflect the developmental stages approaching gastrulation^7, 16, 20, 22–24^. Mimicking stable co-development of embryonic and extra-embryonic cell types through post-implantation embryonic stages from an initial 3D state is technically difficult to control.

**Figure 1:**
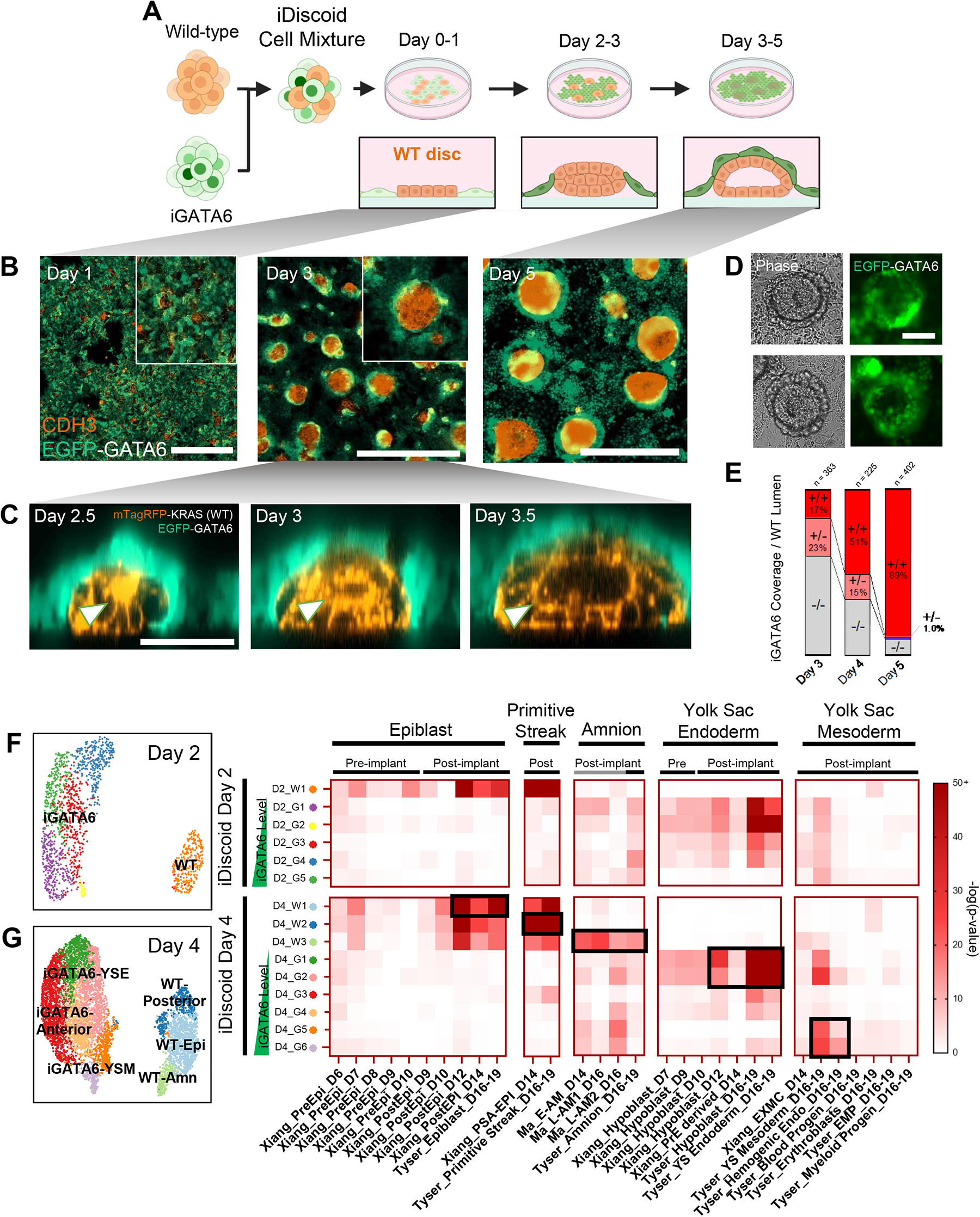
Engineering co-development of embryonic and extra-embryonic endoderm tissues in iDiscoids. (A) Schematic demonstrating cell mixing and subsequent organization of iGATA6 (green) and WT (strongly orange) hiPSCs after GATA6 induction by Dox. (B) Immunofluorescence staining demonstrating self-organization of iDiscoids in the cultures. Scale bar = 1000 µm (C) Live time lapse images of one iDiscoid showing growth of a central lumen from a rosette indicated by an arrow. Scale bar = 50 µm. (D) Phase and fluorescence images from live day 4 iDiscoid cultures showing developed iDiscoid morphologies with EGFP-expressing iGATA6 around a WT cluster. Scale bar = 100 µm (E) Quantification of characteristics of WT clusters possessing different iGATA6 coverage and lumen formation characteristics. No islands without coverage were observed possessing a lumen (-/+ = 0%) (F) Single cell UMAP and hypergeometric statistical comparisons of differentially expressed gene (DEG) lists between day 2 iDiscoid and human and NHP embryo single cell datasets. Labels above heatmaps indicate pre-implantation versus post-implantation embryo sample comparisons; gray bars under labels indicated NHP comparisons. (G) Single cell UMAP and hypergeometric statistical comparisons of differentially expressed gene (DEG) lists between day 4 iDiscoid and human and NHP embryo single cell datasets. Black boxes highlight high DEG similarity (high P values) to relevant populations of interest.

To address these challenges, we aimed to develop an alternative strategy to generate structures off the cell culture plate via igniting 2D-to-3D self-organization of hiPSCs. The iGATA6-hiPSCs were mixed with WT hiPSCs, and the mixed population of cells were seeded onto standard culture plates at a defined ratio and cell density (**Fig. 1A**. After treatment with Dox, the iGATA6 cells expectedly upregulate GATA6 and EGFP and exhibit loss of the pluripotency markers NANOG **(Supplementary Fig. 2A)**. During the first 48 hours, from an initial state of random distribution, induced iGATA6 cells and WT cells proliferate, and WT cells are organized into disc-shaped clusters confined by iGATA6 cells **(Supplementary Video 1, Fig. 1B, Supplementary Fig. 1G)**. Subsequently, the migration of the iGATA6 cells over the upper surface of these WT clusters occurs in conjunction with 3D growth within the WT disc **(Supplementary Fig. 2C, Supplementary Video 2)**. These events result in a membrane of extra-embryonic endoderm-like cells that express GATA4 and SOX17 but lack OCT4 over a multilayer WT structure (**Fig. 1A, 1C-D, Supplementary Fig. 2C-D)**. The process of upward iGATA6 migration is in parallel with the deposition of a laminin membrane surrounding the WT clusters **(Supplementary Fig. 3A-B)**. The laminin deposition by the iGATA6 cells can subsequently trigger cell polarization within cells of the WT cluster, which promotes the conversion of cellular rosettes into lumens (**Fig. 1C, Supplementary Fig. 3C, Supplementary Video 3, 4**)^25, 26^. This is facilitated by expression of PODXL, an apically expressed antiadhesive surface protein implicated in embryonic lumen formation, and ZO-1, an apical tight junction protein **(Supplementary Fig. 3C, D**)^27^. This laminin deposition and lumen formation is not observed in WT-only culture in equivalent conditions **(Supplementary Fig. 3E)**.

We observed that within WT clusters for which an iGATA6 layer was not surrounding the cluster, lumen formation was not observed (**Fig. 1E**). The number of clusters with no coverage by an iGATA6 layer progressively decreases by day 5 as the migration of iGATA6 cells moves forward (**Fig. 1E**). Ultimately, following expansion of the lumen, the two cell monolayers of epiblast-like WT (NANOG^+^) and endoderm-like GATA6^+^ cells are positioned on either side of a laminin membrane (**Fig. 1A, 1C, Supplementary Fig. 3C)**, forming a cellular arrangement similar to the bilaminar disc in the early post-implantation human embryo **(Supplementary Fig. 1E)**^28^. Analysis of iDiscoid culture media supported high levels of secreted AFP and APOA1, two proteins known to be produced by primitive endoderm and yolk sac tissues **(Supplementary Fig. 3F)**.

In our initial analysis, we observed that for WT clusters at day 5 with higher area and often lower circularity, cavity formation occurred via multiple lumens **(Supplementary Fig. 4A, B)**. In order to minimize the occurrence of large WT clusters with multiple lumens and control the cluster size and number, we tested a range of seeding ratio and densities for iGATA6 and WT cells, with the objective to achieve cluster area most consistently within a size range corresponding to the E9-E17 human bilaminar disc **(Supplementary Fig. 4C, D)** (Carnegie #8004, #7700 in ^29^, Carnegie #7801 in ^30^). To this end, we selected a seeding ratio of 81:5 iGATA6:WT at a density of 54k cells per cm^2^ (**Supplementary Fig. 4C, D**, dotted boxes). When we applied this optimized seeding density, we could detect 74% of iDiscoids in the expected physiological size range **(Supplementary Fig. 4E)**. Importantly we also observed the majority (70.9%) of iDiscoids in this size range produced a single lumen **(Supplementary Fig. 4E)**. Hence, by altering the initial parameters we could control iDiscoid size, number, circularity, and lumenogenesis.

### Single cell RNA transcriptomics reveals embryonic and extra-embryonic fate trajectories

To investigate the cellular identities of the developed population, we performed single cell RNA sequencing (scRNA-seq) from day 0 to day 5 of iDiscoid cultures **(Supplementary Fig. 5-7)**. Performing an unsupervised comparison of the transcriptomes with the E16-19 human embryo^31^, E6-E14 human embryo samples^24, 32^, and the E18-20 cynomolgus embryo^33^ revealed the acquisition of distinct post-implantation cellular fates in identified iDiscoid subpopulations (**Fig. 1F, G, Supplementary Fig. 6A-C)**.

Beginning as early as 24 hours (D1) after induction, iDiscoids showed the emergence of cell populations (D1_G1, G2, G3) displaying transcriptomic similarity to human hypoblast, alongside a corresponding epiblast-like population (D1_W1) **(Supplementary Fig. 6A-C)**. By 36 hours (D1.5), iGATA6 cells become statistically similar to the human E12 hypoblast lineage in from Xiang et al., and this similarity strengthens through day 5 of culture **(Supplementary Fig. 6B)**^32^. Comparison with the E16-19 human and cynomolgus embryo shows significant similarity to yolk sac endoderm and hypoblast appearing at D1 and strengthening through day 5 (p = 6.00×10^-54^ to 2.84×10^-93^ similarity significance to YS Endoderm at D5) (**Fig 1F, Supplementary Fig. 6A, 6C)**.

On day 2, there is significant statistical similarity to yolk sac endoderm within the iGATA6 population (Clusters D2_G1, D2_G2, D2_G3, D2_G4; p = 1.3×10^-17^ to 9.5×10^-69^), with the highest GATA6-expressing cells (Cluster D2-G5) diverging from this fate towards a more yolk sac mesoderm-like state (p = 6.8×10^-11^) (**Fig. 1F, Supplementary Fig. 6A-C)**. By day 4, we observed the strengthening of the identities of subpopulations within the iGATA6 and fate divergence within the WT compartment. The lower-GATA6 clusters (D4_G1 and D4_G2) showed increased expression of key yolk sac marker genes (*GATA4*, *PDGFRA*, *LAMA1*, *CUBN*, *AMN*, *NODAL*) and statistical similarity to human yolk sac endoderm (**Fig. 1G, Supplementary Fig. 5B, 6A-C**, p_G1_ = 1.3×10^-92^, p_G2_ = 1.8×10^-63^)^34^. A “medium” GATA6 population (D4_G3 and D4_G4) showed lower statistical similarity to either of these fates, but has a strong anterior hypoblast identity (homologous to mouse Anterior Visceral Endoderm, AVE), with specific upregulation of anterior markers (*LHX1*, *HHEX*, *FZD5*, *CER1*, *LEFTY1*) **(Supplementary Fig. 5B)**^31, 35–37^. The highest-GATA6 clusters, D4_G5 and D4_G6, demonstrated statistical similarity to human yolk sac mesoderm (**Fig. 1G, Supplementary Fig. 6A,B**, p_G5_ = 1.7×10^-22^, p_G6_ = 2.5×10^-25^). Taken together, this demonstrates a divergence in yolk sac-associated fates within iGATA6 cells that is correlated with different GATA6 expression levels, with yolk sac endoderm, anterior hypoblast, and yolk sac mesoderm-like cells stratifying between populations expressing low, medium, and high average levels of *GATA6* **(Supplementary Fig. 6** top row). Recently observations in cynomolgus monkey by Zhai et al. suggest a similar stratification of GATA6 levels in a variety of tissues, notably between yolk sac endoderm (lower GATA6) and mesoderm (higher GATA6) populations^38^.

Within the WT clusters, identified via a lack of *GATA6* or *EGFP* transcripts, we observe the divergence of three separate populations from the epiblast-like compartment. The first and largest cluster (D4_W1) maintained the similarity to human epiblast, with expression of the pluripotency markers *POU5F1* (OCT4), *SOX2*, and *NANOG* **(Supplementary Fig. 5B,** p_W1_ = 1.15×10^-49^). The second (D4_W2) WT cluster had lower statistical similarity to human epiblast (p_W2_ = 3.4×10^-20^ to D16-19 Epi), but stronger similarity to human primitive streak (p_W2_ = 3.6×10^-95^ to D16 PrS) (**Fig. 1G, Supplementary Fig. 5B, 6A)**. Our assessment of the third wild type cluster at this timepoint, (D4_W3) showed occurrence of a unique transcriptomic similarity to human and cynomolgus amnion (p_W3_ = 8.0×10^-15^ and 1.6×10^-25^, respectively) (**Fig. 1G, Supplementary Fig. 5B, 6A, 6C)**^17, 39, 40^. We did not observe a similarity to the trophoblast lineages **(Supplementary Fig. 6)**.

When we integrated all clusters across different days, iGATA6 and WT lineages occupied distinct compartments within the projection space. However, we did observe spatial positioning of populations aligned with their time of sampling **(Supplementary Fig 7A)**. Populations sampled at a different time can show similar fates (i.e., D3 versus D5 iGATA6 yolk sac endoderm cells) **(Supplementary Fig 7B)**. The top upregulated genes show distinct patterns across different cell fates, with overlapping genes present across the similar fates sampled at different time points **(Supplementary Fig 7C)**.

### Specification of amniotic ectoderm in iDiscoids

Primate amniogenesis is structurally and temporally distinct from the process in mouse, highlighting the need for human model systems to provide a window into understanding human-specific developmental dynamics^40^. *In vivo*, following lumenogenesis within the epiblast of the primate post-implantation embryo, the cells of the epiblast undergo dorsal-ventral (D-V) patterning to separate the epiblast from amniotic ectoderm^15, 17, 29^. Our single cell transcriptomic analysis in wild type cells showed a group of cells with transcriptomic similarity to human and cynomolgus amnion that emerges around day 3 (**Fig. 2A-B, Supplementary Fig. 6A, 7B)**. These cells express amnion markers, including *ISL1*, *TFAP2A* (AP-2**α**), and *GATA3*, maintaining expression of OCT4 while substantially lacking expression of *NANOG* (**Fig. 2A-B**)^17, 39, 40^. Our analysis via immunofluorescent staining corroborated this finding, demonstrating a layer of ISL1^+^/AP-2**α**^+^ in iDiscoids (**Fig. 2C-E, Supplementary Fig. 8A)**. We observe that these cells are spatially segregated within the cavitated sac-like structure, with the amnion-like lineage positioning away from the bilaminar iGATA6/WT disc, forming a membrane against the tissue culture dish. WT cells above within the bilaminar iGATA6/WT disc display expression of NANOG (**Fig. 2C-E**).

**Figure 2:**
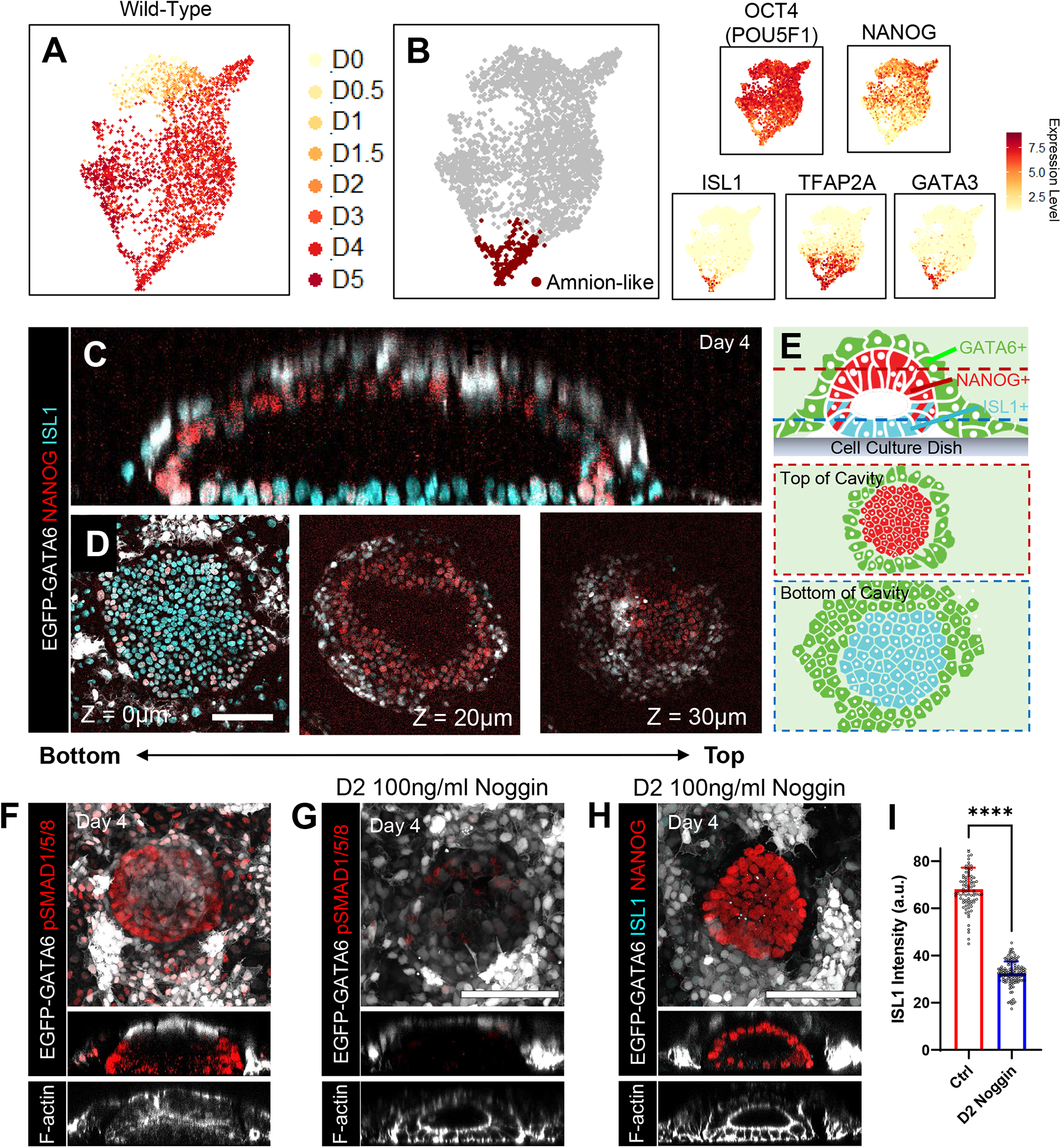
Amniotic-like cavity formation and expansion. (A) Merged UMAP of all WT lineages from iDiscoid labeled by day of development. (B) The merged WT population showing the compartment expressing markers of amnion. *ISL1*, *TFAP2A* (AP-2α), and *GATA3* are expressed in this area, while *NANOG* is negative and OCT4 is low. (C) An orthogonal slice of an individual iDiscoid showing top-bottom compartmentalization of ISL1+ (cyan) and NANOG+ (red) cells while iGATA6 cells cover the top (white). (D) Horizontal z-slices at the indicated distance from the dish of the WT cluster from (C). (E) Schematic showing the position of each population within a single iDiscoid. Dotted lines indicate the area of slices shown. (F) Expression patterns of BMP4 effectors (phosphorylated SMAD1, SMAD5, and SMAD8/9) in iDiscoid. Lower images show a lateral slice of the WT disc shown. (G) Immunofluorescence staining for the BMP4 effectors (phosphorylated SMAD1, SMAD5, and SMAD8/9) in iDiscoid at day 4 after application of the inhibitor Noggin at Day 2 of development. Lower images show a lateral slice of the WT disc shown. (H) Immunofluorescence staining for ISL1 and NANOG in iDiscoid at day 4 after application of Noggin at Day 2 of development. Lower images show a lateral slice of the WT disc shown. (I) Scatterplot showing ISL1 expression (D4) intensities within WT clusters; control (Ctrl) iDiscoid versus BMP4 inhibition (Noggin) conditions. n[control] = 87, n[Noggin] = 134, **** p<0.0001 Scale bars = 100 μm. n represents separate biological samples from a single experiment harvested on the corresponding day.

Further immunofluorescence staining for pSMAD1, pSMAD5, and pSMAD8/9 (BMP4 effectors) revealed BMP4 signal transduction in a ring pattern concentrated at the edges of the iDiscoid (**Fig. 2F**). scRNA-seq at day 4 also showed specific upregulation of BMP4 targets and related genes within the amnion-like population **(Supplementary Fig. 8B)**. Recently, the active role of primate amniotic tissue to promote BMP4 signaling has been suggested, acting downstream of ISL1^40^. It was also reported that BMP4 can promote the formation of ISL1^+^ amniotic tissue in a human embryo model^15, 17, 39^. Hence, we aimed to inhibit BMP signaling via treatment with the inhibitor Noggin to investigate the relevance of amnion-like tissue formation in iDiscoid. We observed effective suppression of SMAD1/5/8 phosphorylation (**Fig. 2G**), and a marked decrease in amnion fate acquisition assessed by ISL1-expression (**Fig. 2H-I**). However, the expression of NANOG remained intact in epiblast-like cells (**Fig. 2H**). To this end, we showed 98.8% of WT clusters express the amnion marker ISL1 around day 4 of iDiscoid development that was consistent across experiments. We show that all (n = 87) but one of the observed WT clusters (n = 88) have ISL1 expression above the expression levels in the noggin treatment group.

Collectively, we conclude that iDiscoids show robust amniogenesis with formation of a dorsal-ventral (D-V) axis that is dependent on BMP4 signaling, supporting the utility of iDiscoids for studying this key developmental event in humans.

### Development of anterior hypoblast poles and posterior axis in iDiscoids

*In vivo* evidence shows a group of cells with anterior visceral endoderm characteristics in human and non-human primates that have been termed anterior hypoblast^36^. In mouse, this tissue has been shown to pattern the specification of the posterior axis via the expression of signaling inhibitors including CER1. However, common *in vitro* stem cell models of the human embryo (i.e., blastoids) have not yet captured this specification^7, 16, 20^. We asked whether iDiscoids can exhibit features associated with anterior-posterior (A-P) axis formation (**Fig. 3**). Our scRNA-seq analysis of iGATA6 cells as early as day 2 revealed a cluster of cells that exhibit markers associated with anterior hypoblast. These cells are positive for *CER1*, *LHX1*, *HHEX*, *LEFTY1*, and *LEFTY2*, and continued to present these markers in the following days (**Fig. 3A**)^31, 35–37^. Additionally, we also detected the emergence of a distinct cluster of cells at day 3 and 4 presenting *TBXT* (Brachyury), *MIXL1*, *GDF3*, and *EOMES*, key markers associated with development of the posterior pole in the embryo (**Fig. 3B**)^41^.

**Figure 3:**
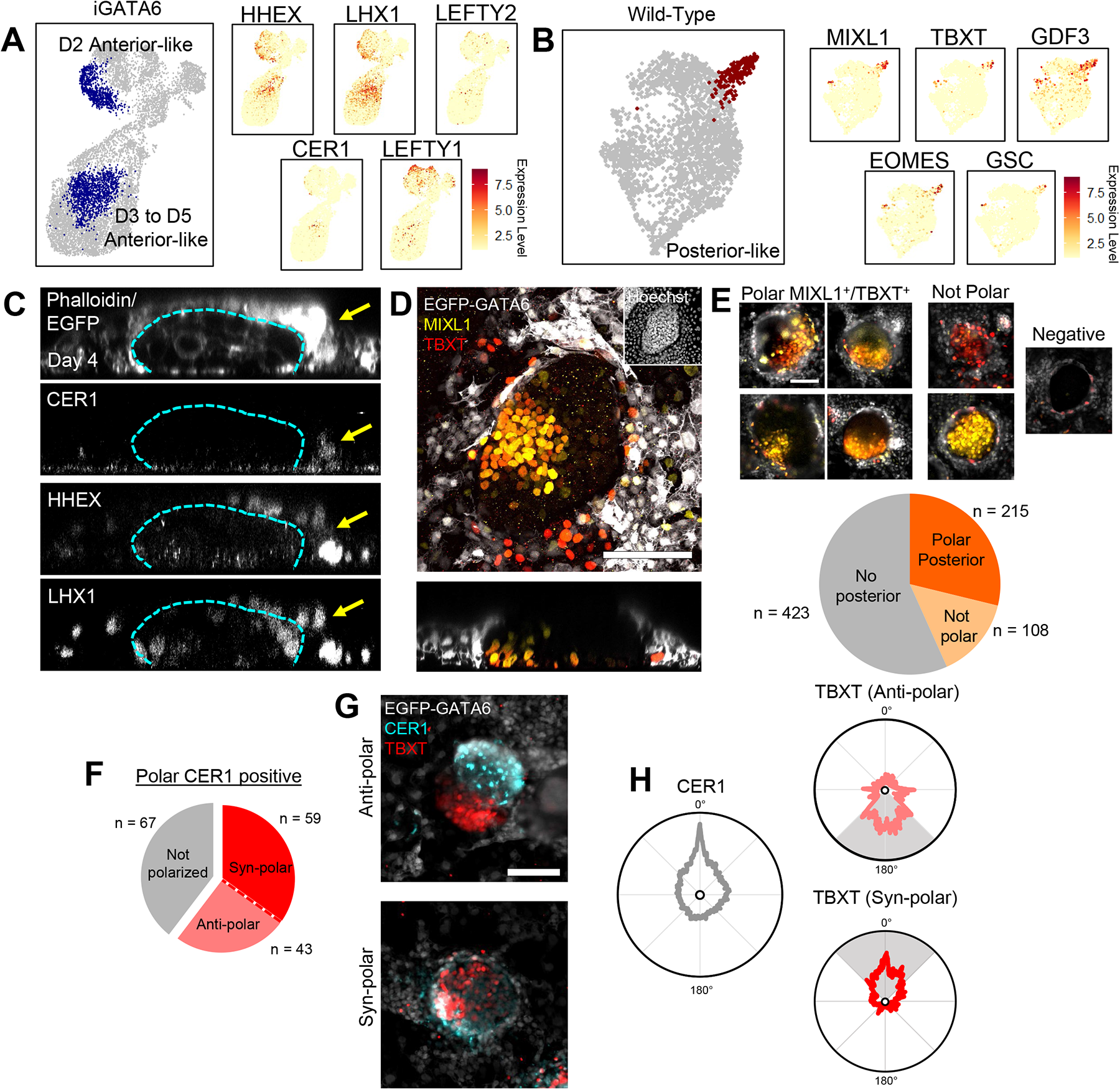
Anterior hypoblast-like domain and posterior axis in iDiscoids. (A) Merged UMAP of all iGATA6 lineages showing the compartments expressing markers of anterior hypoblast. Two separate domains of these markers were observed, one within day 2 iDiscoid cells and one within day 3-5 iDiscoid cells. (UMAP labeled by day can be seen in Supplementary Figure 5) (B) Merged UMAP of all WT lineages showing the compartment expressing markers of the posterior axis and primitive streak. (C) Immunofluorescence staining showing a z-slice of an iDiscoid with a HHEX/LHX1/CER1 co-positive domain adjacent to a WT clusters. Dotted line indicates boundaries of the WT cluster in each image, arrow indicates co-positive domain. (D) Immunofluorescence staining showing a TBXT/MIXL1 co-positive domain within the WT clusters of the iDiscoid. (E) Examples of WT clusters with polarized TBXT/MIXL1 domains (top) or without co-expression of markers (bottom). Venn diagram shows proportions of cluster types observed in iDiscoid cultures, n = 746 total. (F) Venn diagram showing the proportion of clusters with a particular TBXT/CER1 polarity type when CER1 confined to one pole is present. n = 169 total. (G) Representative WT clusters showing TBXT and CER1 expression domains at opposing poles of the cluster (anti-polar, top) or at the same pole of the cluster (syn-polar, bottom). (H) Diagrams indicating the average radial expression patterns of TBXT in each polarity pattern from WT clusters with a polarized CER1-expressing domain indicated in (F). All diagrams are scaled to the same expression intensity value (1 a.u.). Degrees indicate radial distance around the circularized perimeter of a WT cluster from the CER1 peak shown. Shaded areas indicate region of highest average TBXT polarity corresponding to the polarity types. Scale bars = 100 µm. n represents separate biological samples from a 3-4 separate experiments harvested on day 4 after iDiscoid induction.

Immuno-staining identified domains of cells expressing the anterior hypoblast-related proteins (CER1, LHX1, HHEX) in a subset of extra-embryonic cells surrounding the WT clusters (**Fig. 3C**). Out of 396 of iDiscoids assessed we identified 42.4±4.1% (169) discs with polar CER1-expressing domains on day 4 (**Fig. 3C**), exhibiting a similar efficiency to existing *ex vivo* human blastocyst cultures^36^. In parallel, we showed development of TBXT^+^ domains in 46.4±9.0% (323 out of 746) of WT clusters on day 4 in iDiscoids, of which 30.1±6.3% (215) of clusters with these domains displayed asymmetric (polar) expression (**Fig. 3D-E**). These posterior domains are dependent to local BMP4 signaling, as a small molecule, Noggin, could eliminate TBXT^+^ poles with no effect on development of CER1-expressing cells. This supports a role for BMP4 from amnion in posterior axis specification in line with past studies within these tissues in iDiscoids (**Supplementary Fig. 9**)^42^.

We then analyzed iDiscoid cultures to evaluate A-P axis positionings. On day 4, we observed that 60.4±5.3% (102 of 169) of iDiscoids with polar configuration in CER1-expressing cells possessed a TBXT-expressing pole (**Fig. 3F**). Within iDiscoids with polarity in both CER1 and TBXT we observed two distinct patterns (**Fig. 3F, G, Supplementary Fig. 10**). 42.2±3.2% exhibit anti-polar configuration, where the TBXT^+^ pole occupies the opposite radial region as the CER1^+^ domain, similar to A-P axis positioning in mouse at E6.5 (**Fig. 3F-H**). In a second set of polar iDiscoids, we saw high expression of CER1 in the proximity of TBXT^+^ domains (**Fig. 3F-H, Supplementary Fig. 10A**). This set of iDiscoids shows CER1^+^ and TBXT^+^ poles with a same radial region (syn-polar), in contrast to past reported literature. We also noted expression of CER1 in a subset of TBXT^+^ cells in the posterior domains in some cases **(Supplementary Fig. 10A, B)** and subsequent scRNA-seq analysis showed the presence of a subpopulation of *CER1*^+^/*TBXT*^+^ cells in the cluster that represents posterior identity (**Supplementary Fig. 10C**).

Next, we aimed to determine whether CER1 expression adjacent to or within the posterior region observed in iDiscoids can be identified during *in vivo* embryogenesis. To address this question, we first examined the recent spatial scRNA-seq data from human and non-human primates. In marmoset embryos^37^, we noted non-polar *CER1* at CS5, followed by very slight polarization away from the TBXT-expressing posterior at CS6 (subtly anti-polar-like) **(Supplementary Fig. 11A)**. However, in human E16-19 (CS7)^31^, we observed higher *CER1* transcript expression in the same caudal embryonic region as the primitive streak **(Supplementary Fig. 11B)**. Lastly, pseudospatial assessment of the E18-20 cynomolgus embryo shows *CER1* and *TBXT* expression overlapping within the lower region of the embryonic disc (hypoblast) and embryonic disc compartments (syn-polar-like) **(Supplementary Fig. 11C)**^43^. These three datasets show a subset of cells or regions that display expression of *CER1* near or within the posterior domain. To re-examine this phenotype in mouse, we assessed three independent mouse scRNA-seq datasets spanning ∼E6.5-E8 and could also detect a similar pattern of *Cer1* expression within the anterior boundary of the primitive streak **(Supplementary Fig. 11D-F)**^44–46^.

Taken together, our analyses of iDiscoids, in tandem with exploration of available *in vivo* data, suggest that the observed expression and positioning of TBXT^+^ and CER1^+^ cells in iDiscoids can be seen *in vivo* and reveal distinct developmental stages. Anti-polar positioning of CER1 and TBXT domains reflect positioning of anterior hypoblast-like and posterior domains during early primitive streak formation around E14; syn-polar positioning and co-expression of CER1 and TBXT is reflective of a more advanced stages post-E14 where CER1 expression acts either to prevent spreading of posterior domain^47^ or represent early mesendoderm commitment derived from a primitive streak^38^. We propose iDiscoids provide a model to further study A-P axis biology, the formation of anterior hypoblast-like cells, the posterior pole and their respective cellular progeny in a human context.

### Specification of yolk sac mesoderm-like cells with hematopoietic potential

The yolk sac represents a multifunctional extra-embryonic tissue that provides nutritional support for the embryo while also acting as a hub for early blood production that emerges around E16-18 of human development and matures thereafter^2, 48, 49^. Multicellular models of human yolk sac morphogenesis with subsets of mesoderm-derived cells and *in vivo*-like structural characteristics are missing, imposing a challenge for the dissection molecular underpinnings of the process. We detected progressive developmental maturation aligned with yolk sac cellular fates in iGATA6 compartment of iDiscoids (**Fig. 1G, Supplementary Fig 3F, 6)**. We further asked if iGATA6 cells in iDiscoids could mimic development of mesodermal subsets within the developing yolk sac endoderm. Hypergeometric statistical analysis of the clusters with highest expression of GATA6 on day 4 (D4_G6) and 5 (D5_G3) reveals significant similarity to human yolk sac mesoderm, hematopoietic lineage progenitors, and hemogenic endothelial cells as compared with the E16-19 human embryo (**Fig. 4A, Supplementary Fig. 7A)**^31^. Examination of the D5_G3 cluster reveals higher expression of the yolk sac mesoderm marker cell surface marker BST2, as well as of ECM proteins **(Supplementary Fig. 12A)**^49, 50^. Subpopulations of cells within this cluster also exhibit a higher average expression markers of human yolk sac mesoderm (*CREB3L1*, *NR2F2*, *PLAGL1*, *ANXA1*, *NID2*)^31, 40, 50^, markers of endothelial cells (*CD34*, *PECAM1* (CD31), *TEK*, *ICAM1, ETV2*)^31, 51, 52^, and markers of human hematopoiesis (*GATA1*, *GATA2*, *KLF1*, *KLF2*, *ZEB1*, *ZEB2*, *GYPA* (CD235a), and *GYPB* (CD235b)) (**Fig. 4B, Supplementary Fig. 12B)**_31,49,53._

**Figure 4:**
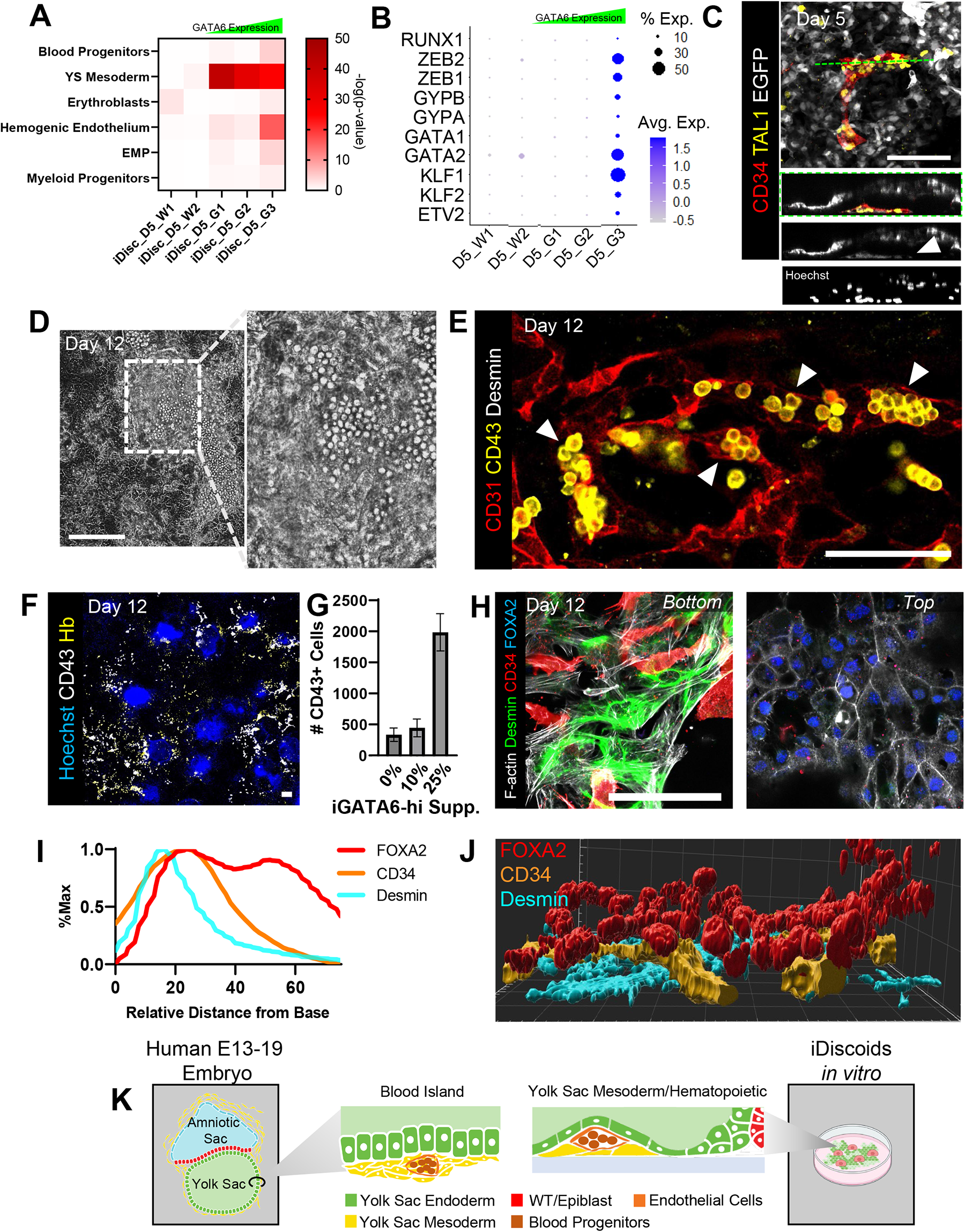
iDiscoid hematopoietic lineages and blood island-like structures. (A) Heatmap showing day 5 iDiscoid scRNA-seq populations compared to hematopoietic populations from the human E16-19 embryo. (B) Dot plot showing the expression pattern of hematopoietic markers in day 5 iDiscoid scRNA-seq populations. (C) Immunofluorescence image showing the distribution of cells expressing CD34 and TAL1 (scl) in iDiscoid culture. Cells expressing TAL1 localize between the yolk sac endoderm compartment and the tissue culture dish and form arrangements of spindle-shaped cells. Orthogonal slice shows the position of spindle cells against the dish. Dashed line indicates the position from which the slice was taken. (D) Live phase image taken at day 12 showing cells spherical cells generated within iDiscoid tissue. Dotted box indicates area of inset. (E) Immunofluorescence image of day 12 iDiscoid showing the generation of CD43+ spherical cells within CD31+ endothelial cells. (F) Immunofluorescence image of a day 12 iDiscoid with 25% GATA6-high cells substituted into the normal iDiscoid ratio. (G) Histogram showing the number of CD43+ cells detected in experiments in which the given percentages of GATA6-hi cells were substituted. n[0%] = 4 experimental replicates, n[10%] = 2 experimental replicates, n[25%] = 2 experimental replicates. (H) Two slices from a single blood island-like structure demonstrating Desmin+ mesoderm cells and CD34+ endothelial cells localized underneath a FOXA2+ endoderm. (I) Histogram showing the z-distribution of the indicated markers between the bottom of the dish and the top of the culture. Bimodal distribution of FOXA2 indicates areas of expression outside of blood island-like foci. Distributions are averaged from 9 foci from 3 technical replicates. (J) 3D reconstruction of one blood island area showing positioning of endoderm (FOXA2), endothelial (CD34), and mesoderm (Desmin) markers (K) Schematic depicting in vivo embryonic yolk sac blood islands and in vitro iDiscoid yolk sac blood island-like structures. Scale bars = 100 µm.

Hematopoietic processes emerge *in vivo* in the form of blood islands within yolk sac tissues. In iDiscoids, immunofluorescence analysis initially reveals spindle shape cells positive for TAL1 (scl), a key regulator of blood development^54^. Further analysis shows a subset of these cells were also positive for CD34 and ERG, markers associated with hematopoietic and endothelial fates or RUNX1, a master transcription factor of hematopoiesis **(Supplementary Fig. 13A-C)**^54^. Further analysis shows a subset of these cells were also positive for CD34 and ERG, markers associated with hematopoietic and endothelial fates or RUNX1, a master transcription factor of hematopoiesis **(Supplementary Fig. 13A-C)**^55, 56^. Image analysis of z confocal sections shows ERG^+^ cells consistently emerged underneath an iGATA6 layer (**Fig. 4C, Supplementary Fig. 13D)**, a morphology that may mimic an initial *in vivo* localization similar to human yolk sac mesoderm, where this tissue is positioned against ECM in the area underneath the yolk sac endoderm^55, 56^. Observed CD34^+^/TAL1^+^ cells were also EGFP^+^, indicating they develop from an iGATA6-derived parental population, supporting a hypoblast origin (**Fig. 4C**). Staining for other endothelial markers KDR (VEGFR2) and CDH5 also confirmed similar localization in day 5 iDiscoids **(Supplementary Fig. 13E)**. A colony-forming unit (CFU) assay initiated with CD34^+^ cells followed by flow cytometry analysis of the cultures, showed generation of CD45^+^ cells, erythroid cells (CD71^+^, CD235a^—^ (early stage) or CD235a^+^ (late stage)) as well as CD11b^+^ myeloid cells **(Supplementary Fig. 13F)**.

Having observed early hematopoietic potential in day 5 iDiscoid, we followed iDiscoids beyond day 5 to better examine the hematopoietic potential of the system. We observed minimal expansion of the CD34+ area when the culture was maintained on mTeSR **(Supplementary Fig. 3G)**. Therefore, to facilitate the differentiation of progenitor cells, we used basal media (IMDM without Dox) after day 5. We observed the expansion of the CD34^+^ endothelial-like cell area significantly between day 5 and day 12, from covering 1.2±0.46% of the culture area on day 5 to covering 14.7±6.0% of the culture area at day 12, an expansion that was also confirmed by flow cytometry analysis **(Supplementary Fig. 13G, H)**. Around this time, we observed the emergence of condensed areas containing spherical cells within the yolk sac-like compartment (**Fig. 4D**). Immunofluorescence staining of these areas revealed the presence of a CD31^+^ endothelial layer surrounding spherical cells expressing Leukosialin (CD43), a marker of hematopoietic progenitors (**Fig. 4E**)^53, 57^. Further investigation of these spherical cells found that many expressed the erythropoietic marker genes CD235a and hemoglobin **(Supplementary Fig. 14A, 15A)**^31^. Having observed yolk sac mesodermal and hemogenic progenitors of this cell type primarily correlating within the GATA6-hi clusters in scRNA-seq, we further supplemented iDiscoid cultures at the time of seeding with high-GATA6 expressing cells and observed significantly increased generation of these CD43^+^ hematopoietic foci (**Fig. 4F-G, Supplementary Fig. 15B)**, demonstrating predictable and controllable fate augmentation at the time of seeding.

Further investigation of the cell types observed at day 12 revealed cells with markers of yolk sac-derived myeloid lineages, including macrophages (CD33^+^/CX_3_CR1^+^) and megakaryocytes (CD41^+^/CD42b^+^) **(Supplementary Fig. 14B-C)**^58–60^. Structural image analysis of CD43^+^ foci show intricate 3D hierarchical organization, with CD34^+^ endothelial vessels packed between FOXA2^+^ endodermal cells occupying the top compartment and Desmin^+^ mesodermal cells forming the basal layer (**Fig. 4H-J, Supplementary Fig. 16A-B**). This is in line with the tissue morphology reported *in vivo* in blood islands of the developing human yolk sac at around E19-23 (**Fig. 4K**)^49^. Late time point analysis of day 21 iDiscoid cultures using scRNA-seq and flow cytometry similarly supports emergence of populations aligned with those expected from the human yolk sac^61^. This analysis shows identification of: *CDH5* (VE-cadherin)^+^ /*CD34*^+^ /*CD31*^+^ /*KDR* (VEGFR2)^+^ /*CD93* (AA4.1)^+^ /*NT5E* (CD73)^—^ /*GYPA*^—^ hemogenic endothelium-like cells; *GYPA*^+^ /*GYPB*^+^ /*TRDN* (CD71)^+^ /*HBG1*^+^ /*HBE1*^+^ /*HBZ*^+^ erythrocyte-like cells; *CD226*^+^ /*CLEC1B*^+^ /*PLEK*^+^ /*GP1BA* (CD42b)^+^ /*GATA1*^+^ megakaryocyte-like cells; and *CD33*^+^ /*CD68*^+^ /*CD45*^+^ /*CD53*^+^ /*CX3CR1*^+^ macrophage-like cells **(Supplementary Fig. 14D-F)**. Flow cytometry analysis of these cultures also corroborated the development of a CD43^+^/CD235a^+^/CD71^+^ erythrocyte-like population, a CD43^+^/CD33^+^/CD31^+^ macrophage-like population, and a CD43^+^/CD42b^+^ megakaryocyte-like population **(Supplementary Fig. 14G)**.

## Discussion

iDiscoids, presented here, develop both embryonic and extra-embryonic lineages with several features observed together for the first time in an *in vitro* stem cell-based human developmental model. These features show close proximity to native post-implantation human embryogenesis yet are uniquely observed together in an hiPSC-based system **(Supplementary Fig. 17)**. The iDiscoid platform shows morphogenesis of bilaminar disc-like structures including the formation of an amniotic cavity, the segregation of amnion from pluripotent epiblast-like cells, the generation of anterior hypoblast-like cells and a posterior axis, as well as yolk sac morphogenesis. Our extended follow up revealed morphologically relevant blood islands and the specification of hematopoietic lineages **(Supplementary Fig. 17)**. In our follow up, WT clusters exhibited characteristics mainly aligned with ectodermal fates, which is a subject for future studies and optimization **(Supplementary Fig. 18)**. We also show the utility of the iDiscoid platform to study developmental processes.

From an engineering angle, iDiscoid platform highlights the power of tissue niche engineering and co-differentiation to promote morphogenetic events *in vitro*. It demonstrates that a combination of genetic circuit-based tissue engineering with the formation of tissue-tissue boundaries enables geometric confinement and subsequent self-organization in 3D. Use of genetic circuits can also alleviate the need for complex regimes of growth factors to support heterotypic tissues. The extra-embryonic differentiation of iDiscoids is pre-programmed into undifferentiated hiPSCs via an inducible genetic switch and mixed with WT hiPSCs; hence, the undifferentiated cell mix can be expanded, cryostored, and shipped for use on demand, requiring only 2D culture plates, commercially available medium, and addition of a small molecule, Dox **(Supplementary Fig. 19)**. The high-throughput format, efficient generation, compatibility with live imaging, and an easy-to-implement protocol enables establishment and utility across different labs. These characteristics, coupled with the potential to be produced from hiPSCs with diverse genetic backgrounds **(Supplementary Fig. 20)**, facilitate the studies on early stages of post-implantation, opening new routes for drug testing, developmental toxicology, and tractable disease modeling with fewer ethical concerns in the human context.

## Methods

### Ethics regarding development of iDiscoids

This research was approved and performed under the oversight of the University of Pittsburgh Human Stem Cell Research Oversight Committee. The work that has been reported in this study followed 2016 Guidelines for Stem Cell Research and Clinical Translation as well as 2021 ISSCR Guidelines on Ethical Standards for Stem Cell Embryo Models^62, 63^. To this end, iDiscoids do not involve human embryonic stem cells and are generated from human iPSCs that are derived from human somatic cells such as fibroblasts. iDiscoids are attached to a cell culture dish, lacking an extra-embryonic trophectoderm tissue critical for full integrated embryo development and implantation in the uterine cavity. The yolk sac cavity in iDiscoid is not closed, and the tissue cannot be harvested for any implantation without substantial disruption of their structures. TBXT^+^ posterior domains were observed during the study but were not sustained during the development of our system. These features collectively restrict the ability of this model to undergo the full integrated development of a human embryo *in vitro* and/or implant to support further development of a conceptus *in vivo*.

### Cell Culture

All cells and tissues were cultured in a humidified incubator at 37°C and 5% CO_2_. Our hiPSC lines were cultivated under sterile conditions in mTeSR-1 (Stem Cell Technologies, Vancouver), changed daily. Tissue culture plates were coated for 1 hour at room temperature with BD ES-qualified Matrigel (BD Biosciences) diluted according to the manufacturer’s instructions in ice cold DMEM/F-12. Routine passaging was performed by incubating hiPSC colonies for 5 minutes in Accutase (Sigma) at 37°C, collecting the suspension and adding 5mL DMEM/F-12 medium containing 10 µM Y-27632, centrifuging at 300g for 5 minutes, and resuspending in DMEM/F-12 supplemented with 10 µM Y-27632 for counting. Cells were seeded at a cell density of 25,000 cells per cm^2^ for routine maintenance.

### GATA6-Engineered Cell Line Generation

rtTA-expressing PGP1 hiPSCs previously generated^64^ were transfected using Lipofectamine 3000 (Thermo Fisher Scientific) with Super PiggyBac Transposase (System Biosciences) and the PiggyBac transposon vector with human GATA6-2A-EGFP under control of the tetracycline responsive element promoter. Transfected cells were selected by adding 0.5mg/mL puromycin to the mTeSR1 maintenance medium. PGP9 iGATA6 hiPSCs were engineered as explained previously^64^. For the generation of GATA6-hi cell line, the iGATA6 cell line was sorted via FACS (one cell per each well of a 96 well plate) in mTeSR-1 supplemented with 10 µM Y-27632 and 2µM Thiazovivin. The media was replaced on the day after sorting with mTeSR-1 supplemented with 10 µM Y-27632 and 2µM Thiazovivin. On day 3 after sort, 125µl of mTeSR-1 was added to each well. The wells were monitored afterwards for colony formation. On day 6, rtTA-expressing PGP1 hiPSCs previously generated^64^ were transfected using Lipofectamine 3000 (Thermo Fisher Scientific) with Super PiggyBac Transposase (System Biosciences) and the PiggyBac transposon vector with hGATA6-2A-EGFP under control of the tetracycline responsive element promoter. Transfected cells were selected by adding 0.5mg/mL puromycin to the mTeSR1 maintenance medium. PGP9 iGATA6 hiPSCs were engineered as explained previously^64^. For the generation of GATA6-hi cell line, the iGATA6 cell line was sorted via FACS (one cell per each well of a 96 well plate) in mTeSR-1 supplemented with 10 µM Y-27632 and 2µM Thiazovivin. The media was replaced one the day after sorting with mTeSR-1 supplemented with 10 µM Y-27632 and 2µM Thiazovivin. On day 3 after sort, 125µl of mTeSR-1 was added to each well. The wells were monitored afterwards for colony formation. On day 6, the wells with a considerable colony were passaged and the level of GATA6-EGFP was characterized by dox induction.

### Generation of iDiscoid

The GATA6-engineered hiPSCs were seeded at a ratio of 81:5 with rtTA expressing PGP1 hiPSCs either containing or lacking an mKate reporter gene at a total density of 54,000 cells per cm^2^ in mTeSR-1 supplemented with 10 µM Y-27632. The next day, the medium was changed to mTeSR-1 with 1 μg/mL doxycycline to induce expression of the GATA6 transgene, and this medium used for daily replacement for up to 5 days. For experiments continued beyond 5 days, medium was switched to IMDM on day 6, and half media changes were subsequently preformed daily for up to 21 days. For the mTeSR condition in Supplementary Figure 13E, medium was instead switched to mTeSR on day 6 and cultures were maintained for 12 days.

### iDiscoid Cryostorage

Uninduced iDiscoid cells were incubated with Dispase at 70% confluence for 10 minutes, or until visible lifting of colony edges. Cells were washed twice with DMEM/F-12, then colonies were manually scraped from the plate. Colonies were centrifuged at 300g for 5 minutes, and then were resuspended in Cryostor 10. Cells were cooled at -80°C for 24 hours prior to transfer to liquid nitrogen for long-term storage. Cells were stored in liquid nitrogen for at least 24 hours prior to defrosting.

### Signaling Pathway Inhibition

BMP4 signaling was inhibited via application of 100ng/ml Noggin into the normal media at day 2 of iDiscoid culture. This concentration of Noggin was supplemented into the media changed on subsequent days.

### Staining on glass coverslips

Cells were grown on Matrigel-coated 8mm diameter circular glass coverslips, on 14mm coverslip bottom dishes (Mattek Corporation), or on 8 Well µ-Slide (ibidi). Cultures were fixed for 10 minutes in 4% paraformaldehyde (Electron Microscopy Sciences) at room temperature. Coverslips were then washed three times with PBS followed by 15 minutes permeabilization with 0.2% Triton X-100 in PBS. Subsequently the coverslips were washed three times in wash buffer (0.05% Tween-20 in PBS) for 5 minutes and blocked for 20 minutes in 200 ml wash buffer plus 5% normal donkey serum (Jackson ImmunoResearch Laboratories). The primary antibodies were diluted in 5% normal donkey serum in PBS and incubated with the tissues 1 hour at room temperature followed by three washes in wash buffer for 5 minutes each. The secondary antibodies were diluted in 5% normal donkey serum in PBS and incubated with the tissues 1 hour at room temperature followed by three washes in wash buffer for 5 minutes each. Afterwards, the 8mm coverslips were mounted on microscopy glass slides using ProLong Glass Antifade (Life Technologies), cured overnight at room temperature and then sealed with nail polish. Coverslips in coverslip bottom dishes or µ-slides were stored in PBS at 4°C for 3D imaging.

### Image Acquisition and Processing

Images were acquired using the EVOS M700 automated scanning microscope, Leica SP8 confocal microscope, Sartorius Incucyte Live Cell Imaging System, or Nikon A1 Confocal microscope and processed using Fiji/ImageJ software (NIH)^65^. Any contrast adjustments were made in individual channels and applied evenly across the whole image in that channel. Contrast and color balance for color images was applied evenly across the whole image. 3D reconstructions were generated using the Nikon A1 confocal microscope to generate z-stacks spanning ∼100um deep into the tissues and using Imaris (Bitplane) to construct a 3D volume from the stacks. Time lapse videos were initially processed using Fiji/ImageJ, and annotations were added using the Adobe Premiere software. For Supplementary Video 4, a video moving through the Z-slices was initially recorded performing this action in the NIS Elements HC software (Nikon), and the video was subsequently cropped and processed.

### Analysis of Wild-Type Cluster Areas and Radial Expression

Tiled whole-coverslip images were cropped to a central circumscribed square (typically 9000μm^2^) image for analysis. Wild-type compartment analysis was performed using an in-house MATLAB (Mathworks) pipeline (built in MATLAB R2022b). In brief, wild-type compartments were detected programmatically via thresholding the nuclear dye, f-actin stain, or mKate marker channel for areas of high signal intensity. Maximum individual compartment area and average cell size were defined manually. Compartments were filtered to remove those with areas close to the defined individual cell area via the following equation:

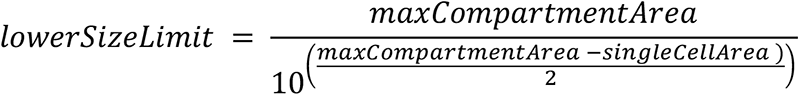

For compartment counting and wild-type area analysis, characteristics of each compartment were recorded using the regionprops function in MATLAB. Additionally, the distance to the nearest wild-type compartment was recorded.

Analysis of marker expression inward from the compartment perimeter was performed via drawing a line a defined distance towards each compartment’s centroid and recording the intensity value of each marker at each point within this range. If a line was drawn such that it would intersect the compartment edge (e.g., drawing from the outer edge of a concave shape), those values were skipped. Intensity values recorded for each line were averaged to create final per-compartment expression distributions. These distributions were aligned based on the point of maximum CER1 expression when present, and the lengths of the distributions were equalized to the length of the longest border recorded using the imresize function in MATLAB.

### Analysis of Wild-Type Cluster Features

Coverage of WT clusters was evaluated via widefield images taken of the entire coverslip. Areas in which EGFP expression was observed inside of WT clusters greater than 50% of the cluster area were counted as being covered. Presence or absence of lumens/cavities was evaluated manually, and was detected via the presence of visible rings of PODXL, ZO-1, or F-actin expression within wild-type compartments.

ISL1 expression within the WT disc was recorded via averaging the radial expression values obtained from the protocol above.

Polarity of TBXT and CER1 domains around WT compartments were assessed in the following way: raw values of CER1 and TBXT produced by radial cluster analysis were normalized to cell density by dividing by the F-actin intensity measured at each point. The average of CER1 and T values at all points within 1/8 of the disc radius in either direction away from a given point around each disc was computed for every point recorded. Positivity for the marker was assessed by comparing the minimum and maximum rolling average values for a difference of greater than 0.1 normalized intensity (a.u.). For islands with positivity, the radial index of the highest averaged quadrant value of CER1 was compared to the radial index of the highest average quadrant value of TBXT: a WT compartment with an average peak of TBXT between 135° and 225° away from the CER1 peak was considered to be anti-polar; a WT compartment with a peak of TBXT within 45° of the CER1 peak was considered to be syn-polar. Polarity type assignments per-island were verified manually by eye.

### Analysis of Hemogenic Endothelial and Hematopoietic Marker Genes

Whole-coverslip immunofluorescence images of each marker were cropped into a square, then divided into equivalent quadrants for ease of analysis. WT cluster masks were created via manual thresholding and creation of a binary image using Fiji/ImageJ. These binary images were then used to remove WT clusters from the images via the Image Calculator function applied to the individual channel images. Resulting images were evaluated via using a custom pipeline in CellProfiler^66^. Scatterplots shown were created using the Mean Object Intensity values for each detected cell. CD43 bar graphs were produced via counts of the number of objects detected by the IdentifyPrimaryObjects function applied to the CD43 channel in isolation. Resulting images were evaluated via using a custom pipeline in CellProfiler^66^. Scatterplots shown were created using the Mean Object Intensity values for each detected cell. CD43 bar graphs were produced via counts of the number of objects detected by the IdentifyPrimaryObjects function applied to the CD43 channel in isolation.

CD34^+^ area was assessed via Fiji/ImageJ. The cropping and WT cluster masking steps above were followed. Subsequently, the Threshold command was used to select areas positive for CD34, and thresholded area was assessed via the Measure command. The area of the threshold was then divided by the total cropped image area to determine percent CD34 coverage for each image.

### Quantitative Analysis of Z-Distribution of Blood Island-related Markers

Confocal z-stack images were captured near to the center of each identified area. For each stack, the bottom was defined as the lowest z-index where an in-focus marker could be identified. The sum of the pixel intensity values from all pixels within each slice above the bottom index were summed, then divided by pixel number. The resulting values at each corresponding z-index were then averaged between all samples identified; these values were then converted to a percentage of the max value to produce the graphs shown.

### Enzyme-linked Immunosorbent Assays (ELISA)

Samples were assayed for AFP and APOA1 using commercially available ELISA kits (abcam). Sample dilutions were optimized to attain detection in the linear range of the standard curves for each individual assay.

### 10x Genomics Sample Preparation for Next-generation Sequencing

Samples were prepared as described by the 10x Genomics Cell Multiplexing Oligo Labeling for Single Cell RNA Sequencing Protocols for cells with >80% viability. iDiscoids of each day indicated were acquired in single cell suspension by incubation with Accutase for 20 minutes at 37°C. The cell suspension was passed through a 40μm strainer in order to remove aggregates. Each sample suspension was then centrifuged at 300g for 5 minutes at room temperature. The supernatant was manually aspirated from each of the samples using a P1000 pipette. The cell pellets were each resuspended in 1ml PBS+0.04% BSA, and the samples were centrifuged at 300g for 5 minutes at room temperature. The samples were then resuspended, targeting 1×10^6^ cells/ml based on expected densities of cells at each day, and the cell suspension was passed through a 40um strainer again. Each cell suspension was counted using a hemocytometer, and a volume of cell suspension was removed as necessary to adjust the final cell count to 1×10^6^ cells. The volumes were then adjusted to 1ml by adding additional PBS+0.04% BSA to replace the removed volumes.

#### For Multiplexing Oligo Labeling

Samples were then centrifuged at 300g for 5 mins at room temperature. Supernatant was aspirated manually with a P1000 pipette. Samples were resuspended in 50μl of CellPlex Multiplexing Solution (10x Genomics), with unique multiplexing oligo solutions assigned to each sample. A maximum of 12 samples were labeled in parallel. Samples were incubated for 5 minutes, starting after the oligo solution was added to the last sample.

Following labeling, 1.95ml 1x PBS + 1% BSA was added to each sample, and the solution was mixed thoroughly. Each sample was centrifuged at 300g for 5 minutes at 4°C. The supernatant was manually aspirated with a P1000 pipette, taking care to leave less than 10μl of supernatant remaining in each sample when possible. Tthe samples were resuspended in 2ml 1x PBS + 1% BSA and were mixed thoroughly to wash. This wash and centrifugation step was repeated two additional times; at the last resuspension, enough wash buffer was added to reach a cell count of 1×10^6^ cells/ml, assuming 50% cell loss from the count at the end of the first protocol. Labeled cell suspensions were then put on ice for transfer to the Pitt Single Cell Core for library creation.

Following a final counting and viability analysis, cells and 10x Genomics reagents were loaded into the single cell cassette, with a target of 25000 single cells for analysis, accounting for predicted cell loss and doublets resulting from multiplexing as laid out in the user guide for the Chromium Single-Cell 3’ Reagent Kit (10x Genomics). After generation of GEMs, the cDNA library was prepared by Pitt Single Cell Core staff following the appropriate steps determined by the 10x Genomics user guide. Libraries were sent to the UPMC Genome Center for sequencing by a NovaSeq S4-200 for an intended read depth of 100,000 reads per cell with 150 bp paired end reads. Our downstream analysis from the sequencing data yielded between 40k-150k mean reads per cell in different samples.

### scRNA-seq Sample Processing and QC

The 10x Genomics CellRanger pipeline was used to align reads to the reference genome (GRCh38.84) appended with transgene sequences, to assign reads to individual cells, and to estimate gene expression based on UMI counts^67^.

Single cell data was excluded based on high mitochondrial genome transcript ratio and either high or low feature or UMI counts. Genes with UMI counts in fewer than 5 cells were removed from consideration. For scRNA-Seq data processing and cluster analysis using Seurat^68, 69^, we used the following general standardized pipeline for processing of the Cell Ranger output: SCTransform regressing percent mitochondrial genes, principal component analysis (PCA), and clustering. Jackstraw plots and permuted p-values were used to assist in determining the optimal number of principal components (PCs) needed to summarize the datasets without losing a significant amount of variation. The quality of a range of clustering resolution values was assessed using enrichment of cluster marker genes (genes differentially upregulated in a given cluster relative to all other clusters) with embryo cell type-specific genes. As a quality check, PC and resolution metrics were modulated to yield fewer or additional clusters to confirm that chosen parameters resulted in the most biologically relevant clustering. Visualization was achieved by the use of UMAP plots identifying cells, clusters, and selected gene expression in each cell, as well as heatmaps and violin plots showing the expression level of genes by cluster. Subclustering the WT in the D3 dataset was achieved by subsetting the WT cluster, then computing the distance between rows of the data matrix using Euclidean measurement (dist function in R). Hierarchical clustering was applied using this distance matrix (hclust in R), and used to create a dendrogram of the distance explained by different numbers of subclusters. The number of subclusters was selected based on the highest level of distance observed and applied to the data (clustercut in R). The cluster was then added back to the rest of the dataset and markers were determined.

### Statistical analysis

#### Similarity quantification between lists of marker genes

We quantify the similarity between two lists of marker genes A and B by using the Hypergeometric test for overrepresentation. This is equivalent to a one-tailed Fisher’s Exact Test.

#### Statistical Tests for Bar Graphs

The statistical test used to calculate p-value for the bar graph in Figure 4I was an unpaired, two-tailed t-test. The statistical test used to calculate p-value for the bar graph in Supplementary Figure 13G is a one-way ANOVA with Dunnett’s multiple comparisons test.

### *in vitro* Colony-Forming Unit Assays

Cells were dissociated into single cell solution using accutase (StemCell Technologies). CD34^+^ cells were purified from a portion of the cells using the CD34 Microbead Kit Ultrapure and LS columns (Miltenyi Biotec) according to the manufacturer’s instructions. Colony forming unit assays were performed using methylcellulose containing media (MethoCult H4434 Enriched and MethoCult SF H4636, StemCell Technologies) according to the manufacturer’s instructions. After 14 days in culture, cells were analyzed immunophenotypically via flow cytometry.

### Flow Cytometry Analysis

Input and output cells to the colony forming unit assays were analyzed immunophenotypically via flow cytometry. Cells were stained with human anti-CD235a, anti-CD71, anti-CD45, anti-CD11b, and anti-CD34 antibodies (Biolegend) for 30 minutes on ice. Cells were analyzed using an LSR II flow cytometer (BD Biosciences) using 7-AAD (ThermoFisher Scientific) as a dead cell stain. Data analysis was performed using FlowJo software. An example of gating strategy can be found in Supplementary Figure 21.

To prepare samples, iDiscoids were developed in 4-6 wells of a 6 well plate, and samples were pooled prior to CD34^+^ cell isolation using the MACS separation system (Stemcell Tech).

For flow cytometry analysis of CD34^+^ cells on D5 and D12, iDiscoids were seeded on three wells of a 12 well plate. On the day of harvest, accutase was used to detach the cells. Cells were stained with anti-CD34 (APC) (Biolegend) at the concentration of 1:600 for 30 minutes on ice. Cells were analyzed using an LSR II flow cytometer (BD Biosciences) using 7-AAD (ThermoFisher Scientific) as a dead stain.

## Data Availability

The datasets generated and/or analyzed during the current study are available from the corresponding author on reasonable request.

## Code Availability

All scripts used have been deposited at https://github.com/AmirAlavi/GATA6_R and are available from the corresponding author upon request.

## Acknowledgements

This work was supported by a Startup Fund from the Department of Pathology at University of Pittsburgh. We acknowledge support from the Center for Biological Imaging and the University of Pittsburgh Flow Cytometry Core facilities. Characterization of hematopoietic cellular subsets was partly supported by an R01 from the National Heart Lung and Blood Institute (HL141805) to M.R.E. Development and characterization of the iPSC lines used in these studies were partly supported by an R01 from the National Institute of Biomedical Imaging and Bioengineering (EB028532) and NSF award # 2134999 to M.R.E. The computational analysis of published embryo datasets was partly supported by NIH grants 1R01GM122096, OT2OD026682, 1U54AG075931 and 1U24CA268108 to Z.B.J. This work is also partly supported by the Pittsburgh Liver Research Center (NIHNIDDK P30DK120531), the Cellular Approaches to Tissue Engineering and Regenerative Medicine Training Grant (T32 EB001026), and NIH 1S10OD019973-01. We would like to thank Katherine Helfrich and Michael Calderon at the Center for Biological Imaging for their assistance related to imaging. We also thank Alaina Sutherland, Chloe Hislop, and Raini Porterfield for helpful discussion during this work. Biorender was used to create several of the schematics shown.

## Author Contributions

J.H. and M.R.E. conceived the study; M.R.E., Z.B.J., B.S., and S.K. supervised the project; J.H., M.R.E., S.K., A.A., K.K.F., J.V., R.L., D.S., Z.B.J. conceived the methodology for experiments; J.H., R.S., K.K.F., J.V., R.L., M.N.T., and M.R. performed iDiscoid experiments; J.H., R.S., K.K.F., T.M., M.N.T., S.W. and D.S. performed imaging and analysis; J.H., A.A., and Q.S. performed the bioinformatic analysis of scRNA-seq datasets; R.L. and K.K.F. performed flow cytometry and analysis; J.H., A.A., K.K.F., J.V., R.L., D.S., M.R.E. performed visualization of the data; M.R.E., S.K., Z.B.J., and J.H. acquired funding for the project; J.H. and M.R.E. wrote the original draft of the manuscript; J.H., R.S., K.K.F., M.R.E., B.S., S.M.C.S.L., A.A., J.V., S.K., and Z.B.J. were involved in editing and input to the final draft of the manuscript.

## Extended Data Figures

**Extended Data Figure 1:**
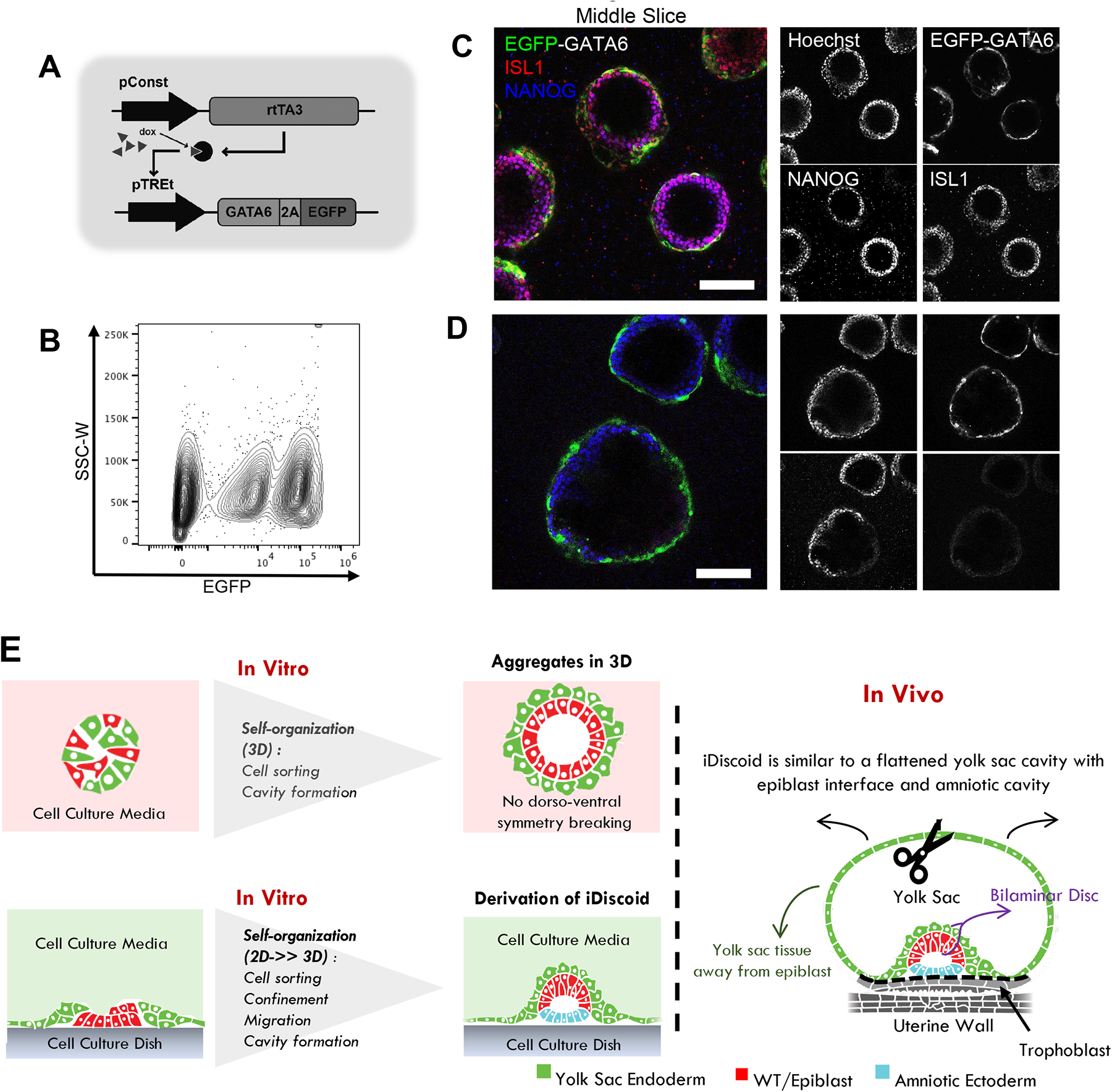
3D iDiscoid cultures. (A) The gene circuit used to create inducible GATA6-expressing iPSCs. pConst is a constitutively active promoter. (B) Heterogeneity of EGFP (GATA6) activation in iGATA6 cells, detected via flow cytometry analysis. Higher gene circuit copy numbers leads to higher expression level of EGFP and GATA6. (C) 3D culture of iGATA4 and WT showing co-expression of the amnion marker ISL1 and the pluripotency marker NANOG spread fully throughout the WT layer without D-V polarization. Middle slice shows the development of a central lumen. (D) 3D culture of iGATA4 and WT showing expression of the pluripotency marker NANOG throughout the WT layer but lack of ISL1 expression in notable subset of these 3D tissues. These spheres do not exhibit polarization as well. (E) Schematic depicting iDiscoid development in 3D versus from 2D>>3D in comparison to embryo morphology. iDiscoid is similar to a flattened yolk sac cavity with epiblast interface and amniotic cavity.

**Extended Data Figure 2:**
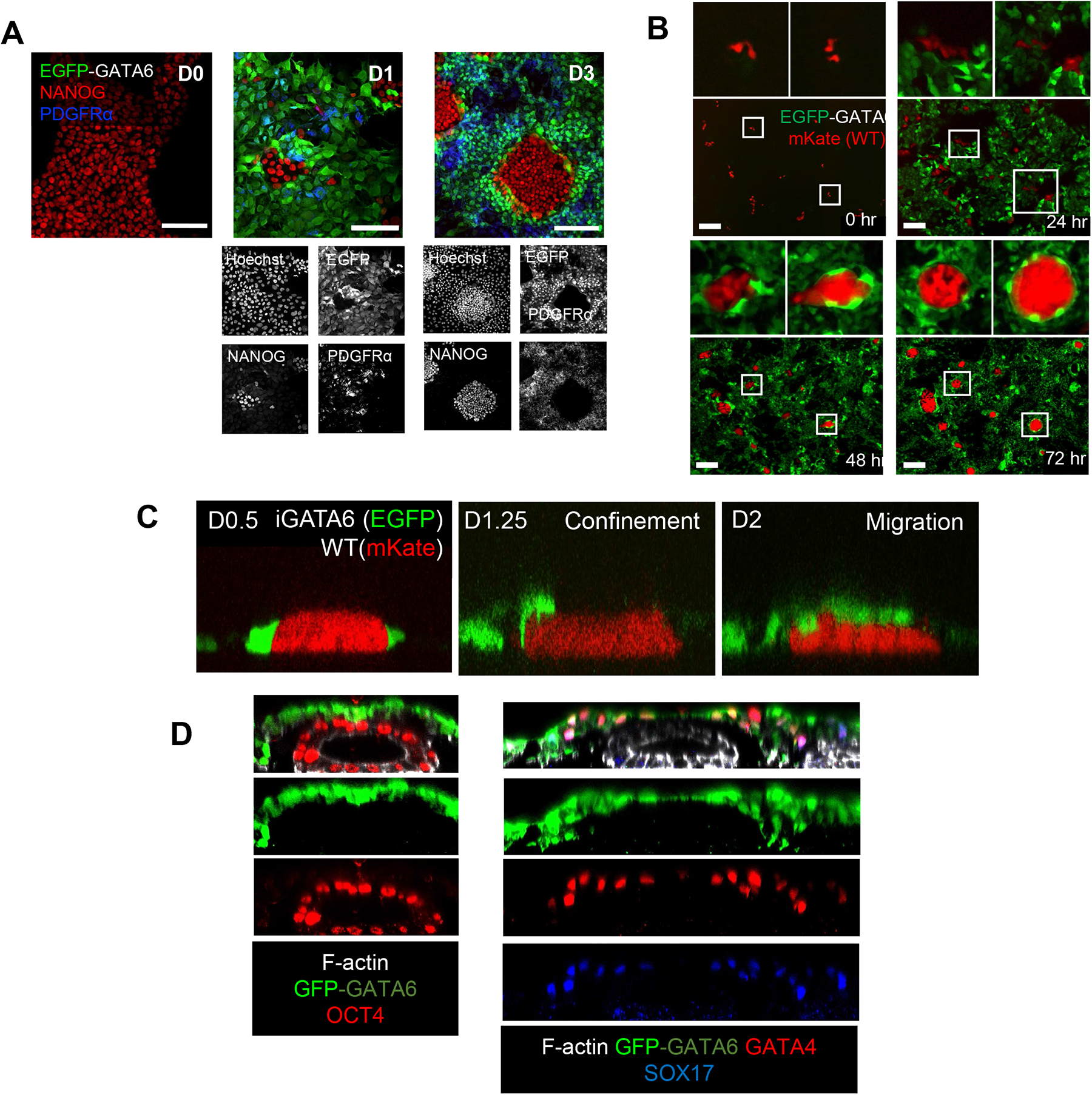
Sorting and symmetry breaking events following GATA6 induction. (A) Immunofluorescence images of fixed cultures demonstrating cell organization of iGATA6 (green) and WT (NANOG) hiPSCs between day 0 and day 3 after GATA6 induction. PDGFRα rises within the iGATA6 cells as they acquire a more yolk sac endoderm-like morphology. (B) Time lapse images of a single position within the iGATA6/WT co-culture from day 0 to day 3 after GATA6 induction. Top cropped images correspond to positions within the white boxes in the images below. Scale bar = 200 μm. (C) Time lapse images showing the initial confinement of red WT cells with green iGATA6 cells, followed by migration of iGATA6 cells over the WT disc prior to D2 after iDiscoid induction. (D) Z-slices of two representative iDiscoids showing localization of OCT4, GATA4, and SOX17 within the bilaminar disc-like area of iDiscoid culture, as well as development of a central lumen by day 5.

**Extended Data Figure 3:**
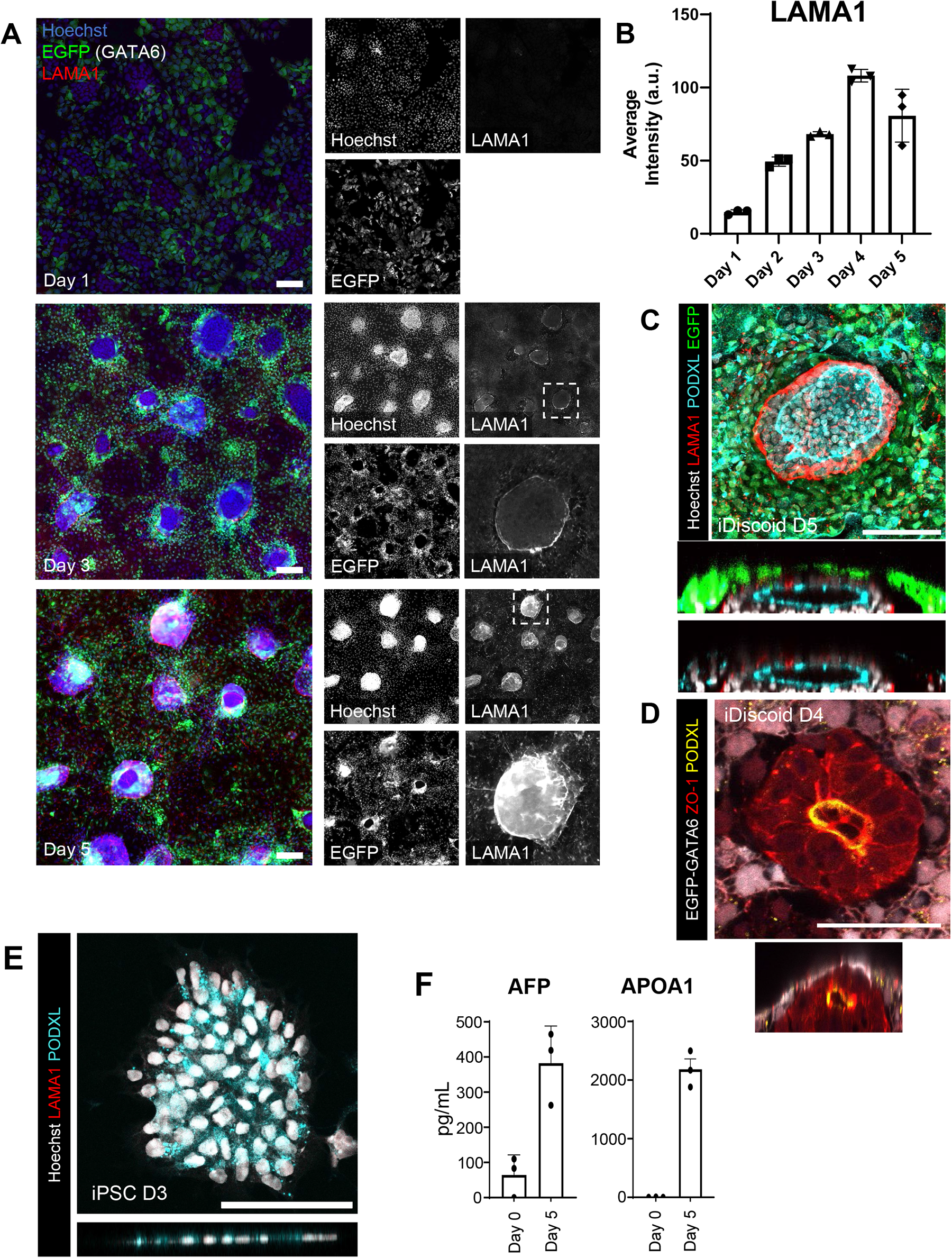
Lumen development within WT of iDiscoids. (A) Immunofluorescence images showing dynamic of LAMA1 deposition on days 1, 3, and 5 of iDiscoid development after induction. Dotted boxes show the areas of inset in the fourth panels of the day 3 and day 5 images. (B) Timecourse graphs showing the increase in LAMA1 signal in immunofluorescence images over the first five days of iDiscoid development. Dots represent randomly sampled areas from a single experiment harvested at each day (C) Immunofluorescence image showing the deposition of laminin around a WT cluster with a central lumen as well as polarization of PODXL . (D) Immunofluorescence showing horizontal and lateral slices of a representative WT cluster with polarization of PODXL and ZO-1 towards a central lumen. (E) A representative cluster of WT iPSCs at day 3. No laminin deposition is observable in the vicinity of the cluster. (F) ELISA comparing secreted AFP and APOA1 detected on D0 and D5 after GATA6 induction with Dox. Individual dots represent biological replicates. Scale bars = 100μm

**Extended Data Figure 4:**
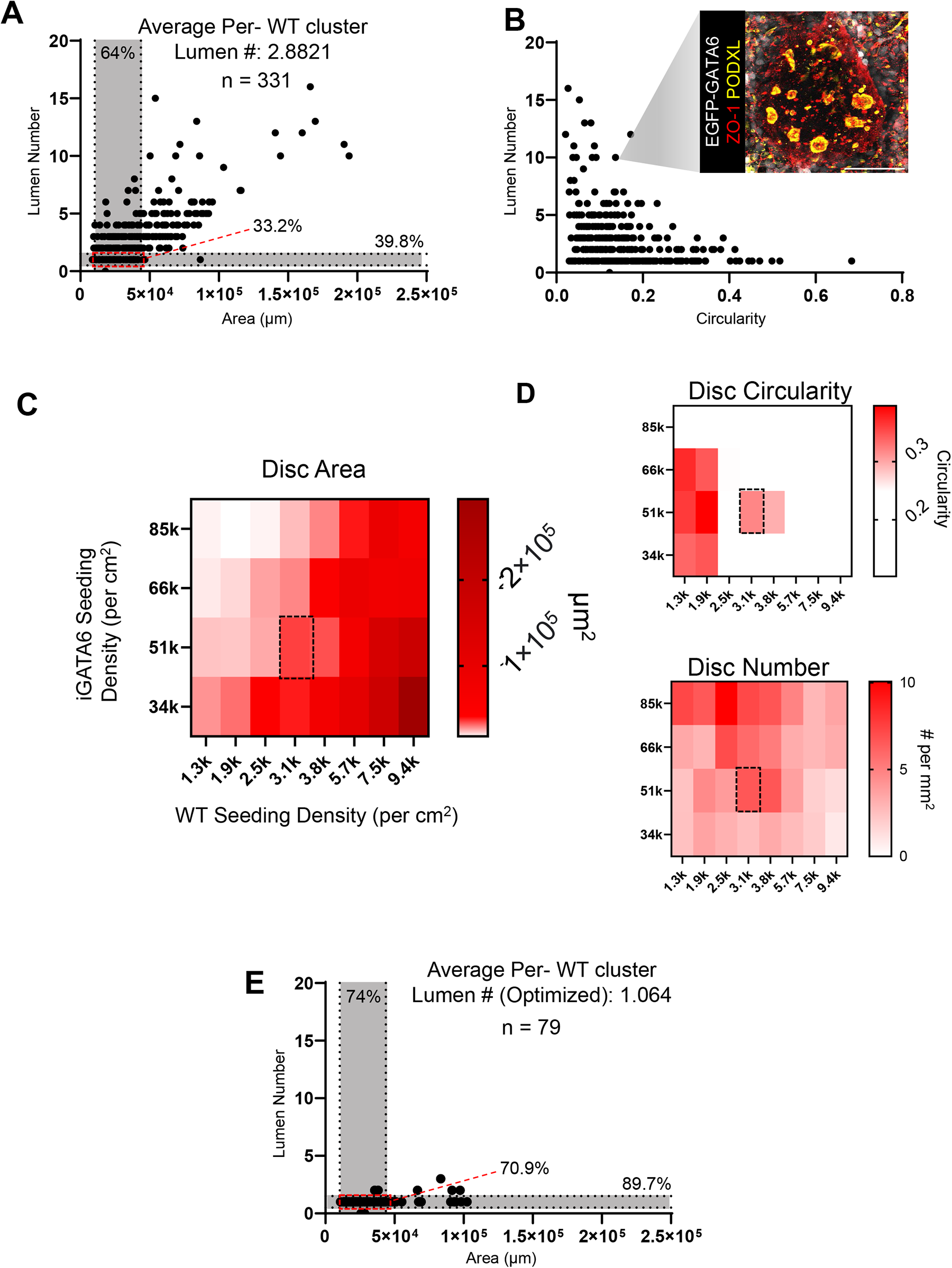
Lumen development within WT of iDiscoids. (A) Distribution of WT cluster areas versus the number of lumens observed in iDiscoids with iGATA6 coverage. Shaded area indicates the areas of the bilaminar disc between E9 and E17 as recorded by Hertig and Rock, 1949 and Heuser et al. 1945; 64% of discs observed have areas that fall into this range of values, and 39.8% have a single lumen. 33.2% fall into both of these categories. n = 331 total. (B) Distribution of WT cluster circularity versus the number of lumens observed in iDiscoids with iGATA6 coverage, with a representative image of an iDiscoid with the indicated characteristics. N = 331, Scale bar = 100μm (C) Heatmaps displaying the average area of iDiscoids resulting from different initial seeding densities of iGATA6 and WT. Dotted box shows optimized seeding density used for iDiscoid experiments. (D) Heatmaps displaying the average circularity and average resulting disc number pre mm2 of iDiscoids resulting from different initial seeding densities of iGATA6 and WT. Dotted box shows optimized seeding density used for iDiscoid experiments. (E) Distribution of WT cluster areas versus the number of lumens observed in iDiscoids with iGATA6 coverage when seeded at the optimized ratio indicated in C. Shaded area indicates the areas of the bilaminar disc between E9 and E16-19 as recorded by Hertig and Rock, 1949 and Heuser et al. 1945; 74% of discs that fall into this area range. 89.7% of discs have a single lumen. 70.9% of discs fall into both of these categories. n = 79 total.

**Extended Data Figure 5:**
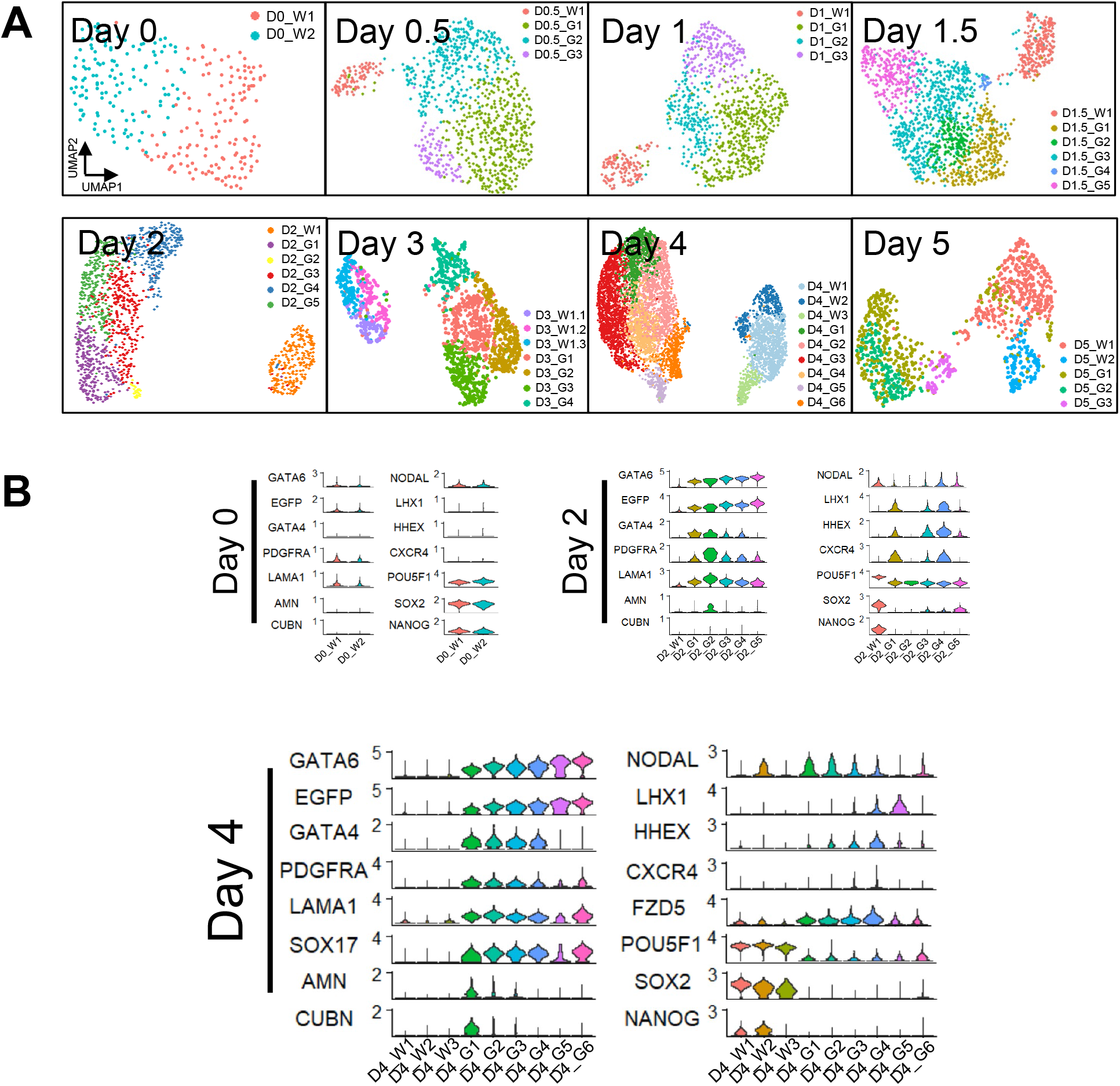
Merged clustering of D0-D5 iDiscoid single cell populations. (A) Individual UMAP projections and clustering for each time point recorded through day 5 in iDiscoid development. (B) Violin plots showing a curated set of genes in iDiscoid day 0, day 2, and day 4 clusters. iDiscoid clusters are ordered by lowest to highest GATA6 expression level. “W” clusters are clusters with putative wild-type lineage; “G” clusters are clusters with putative iGATA6 lineage.

**Extended Data Figure 6:**
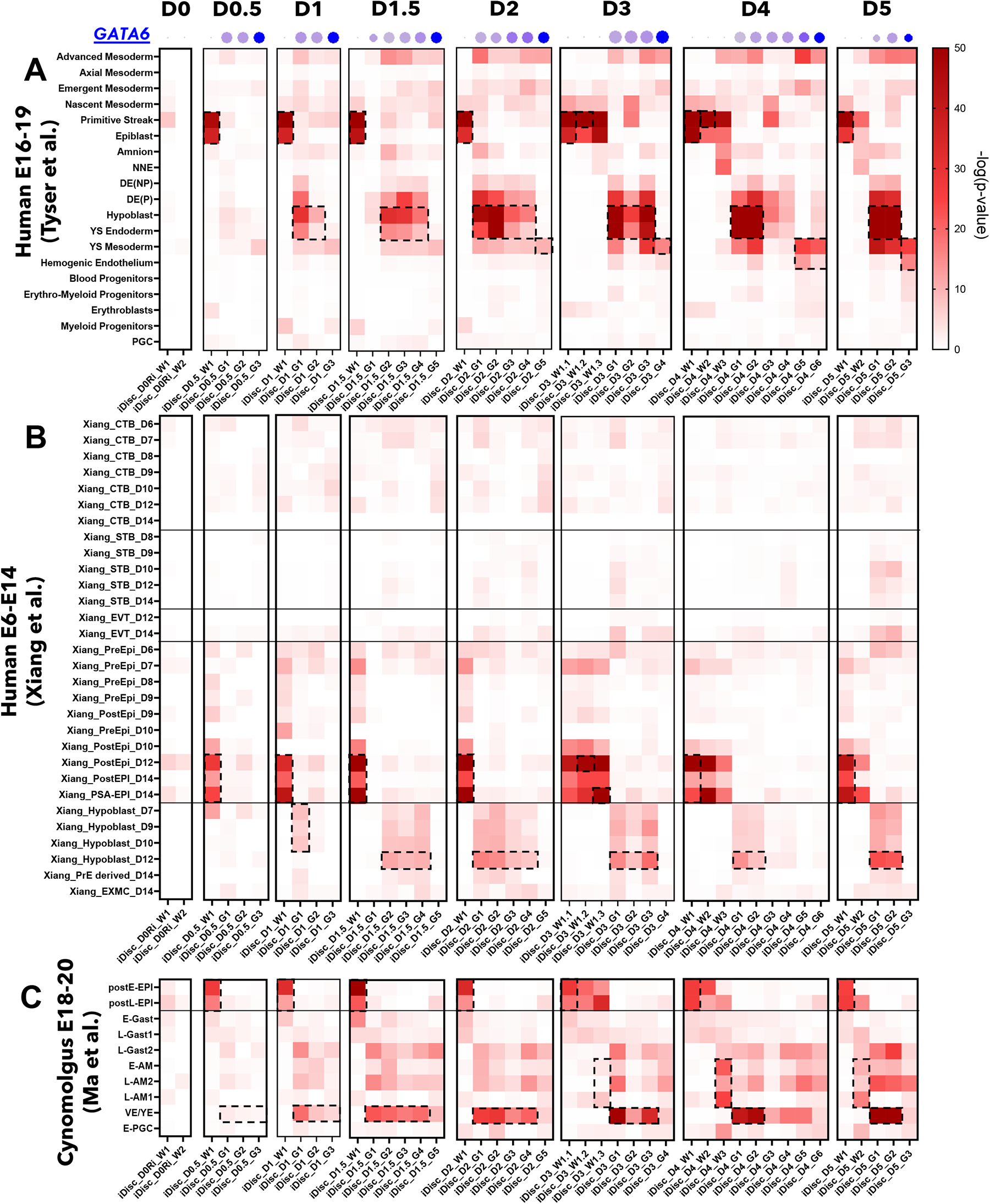
Hypergeometric statistical comparison of iDiscoid time points to human and NHP embryo data. (A) Hypergeometric statistical comparison of each iDiscoid day to the annotated human embryo populations from Tyser et al. 2022. Blue dots above each column indicates the relative GATA6 expression level of each population indicated per day. (B) Hypergeometric statistical comparison of each iDiscoid day to the annotated human embryo populations from Xiang et al. 2020. Scale used is the same as shown in panel A. (C) Hypergeometric statistical comparison of each iDiscoid day to the annotated cynomolgus embryo populations from Ma et al. 2019. Scale used is the same as shown in panel A. iDiscoid clusters correspond to those in the individual day-by-day clustering in Extended Data Figure 5 and are ordered from left to right on the x-axis by lowest to highest GATA6 expression level. “W” clusters are clusters with putative wild-type lineage; “G” clusters are clusters with putative iGATA6 lineage. Dotted outlines indicate fate comparisons of the most interest for each cluster. Abbreviations from other datasets: NNE = Non-neural ectoderm, DE (P) = Definitive endoderm (proliferative), DE (NP) = Definitive endoderm (not proliferative), YS = Yolk sac, PGC = Primordial germ cell, CTB = Cytotrophoblast, STB = Syncytiotrophoblast, EVT = Extravillous trophoblasts, PSA-EPI = Primitive streak anlage in the epiblast, EXMC = Extraembryonic mesoderm cells, E-= Early, L-= Late, Gast = Gastrulating cells, AM = Amnion, VE/YE = Visceral endoderm/Yolk sac endoderm

**Extended Data Figure 7:**
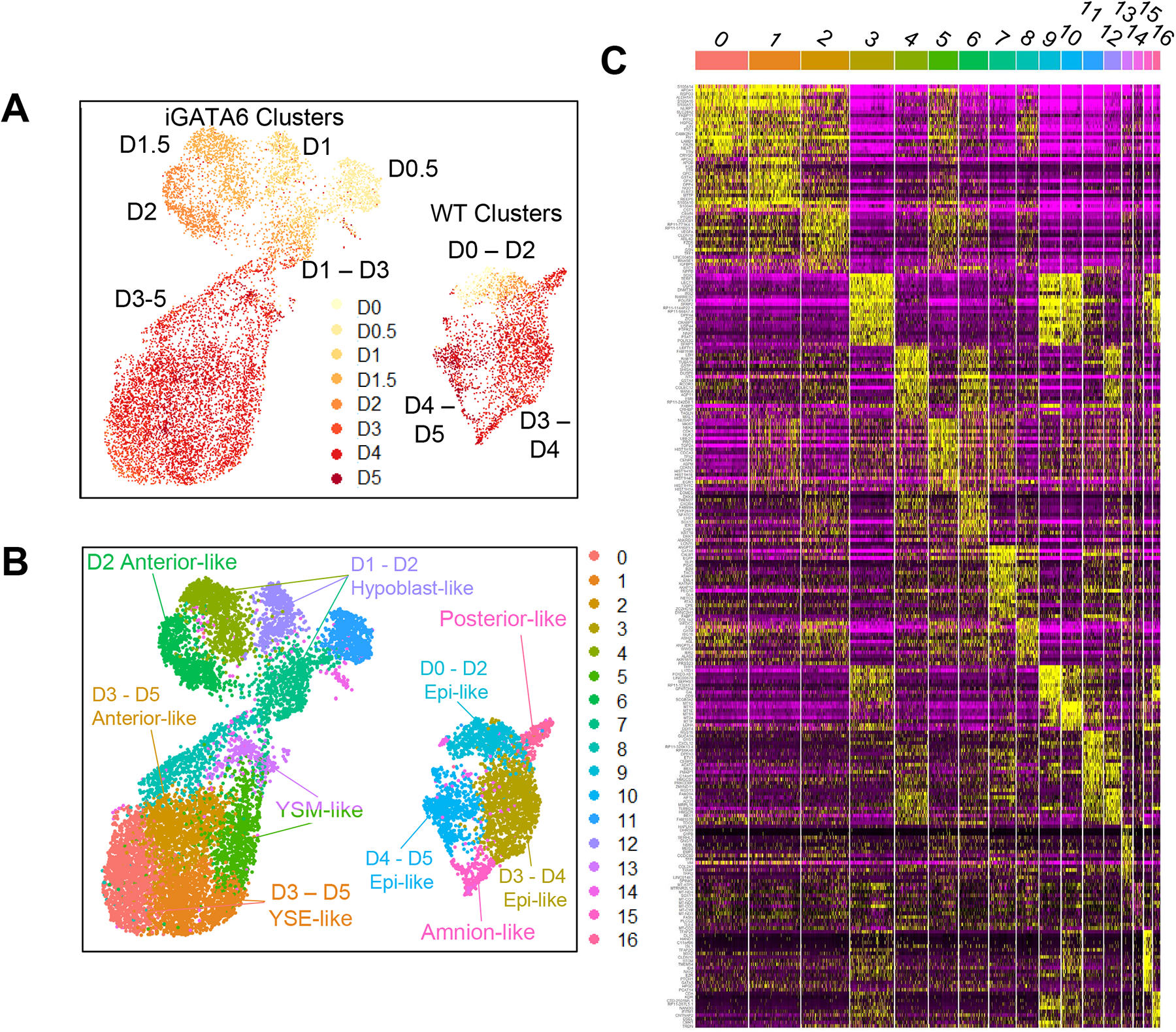
Merged clustering of Day 0 – Day 5 iDiscoid single cell populations. (A) UMAP showing the merged iDiscoid clusters annotated by day. (B) UMAP showing the merged iDiscoid clusters with unsupervised clustering applied. (*repeated figures*: Cluster 14 is the amnion-like cluster shown in Figure 2B; Cluster 6 is the D2 anterior-like cluster shown in Figure 3A; Cluster 2 is the D3-5 anterior-like cluster shown in Figure 3A; and Cluster 15 is the posterior-like cluster shown in Figure 3B.) (C) Heatmap showing the top 20 genes corresponding to each cluster of the merged day 0 through day 5 dataset.

**Extended Data Figure 8:**
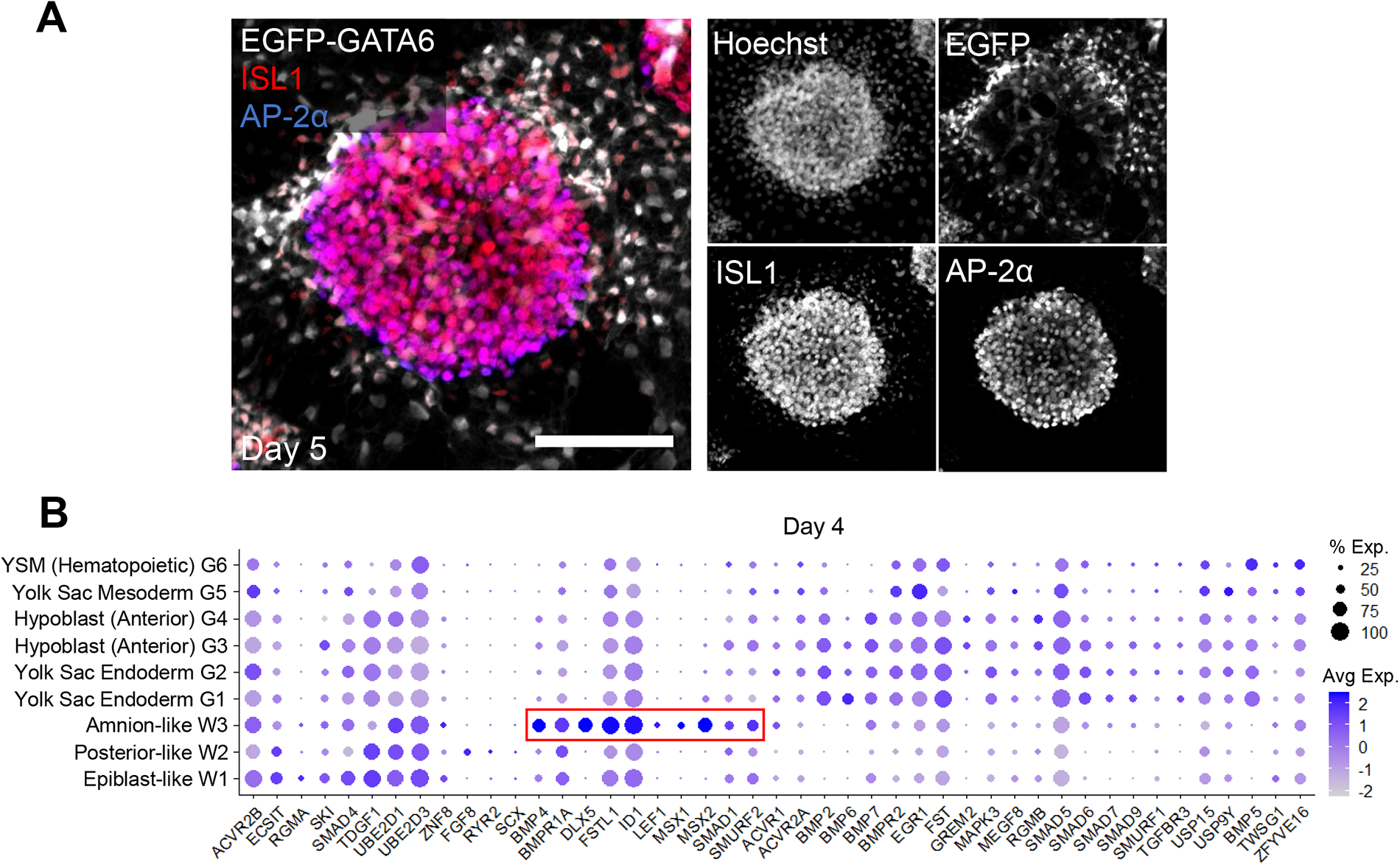
Major signaling interactions between day 5 iDiscoid cell clusters. (A) Immunofluorescence staining for the amnion markers ISL1 and AP-2α at day 4. Top-down widefield image of a flattened coverslip. Scale Bar = 100 μm (B) Dot plot of marker genes from the BMP pathway from day 4 iDiscoid scRNA-seq. BMP4 expression and a number of associated genes (boxed in red) are highest in the amnion-like population.

**Extended Data Figure 9:**
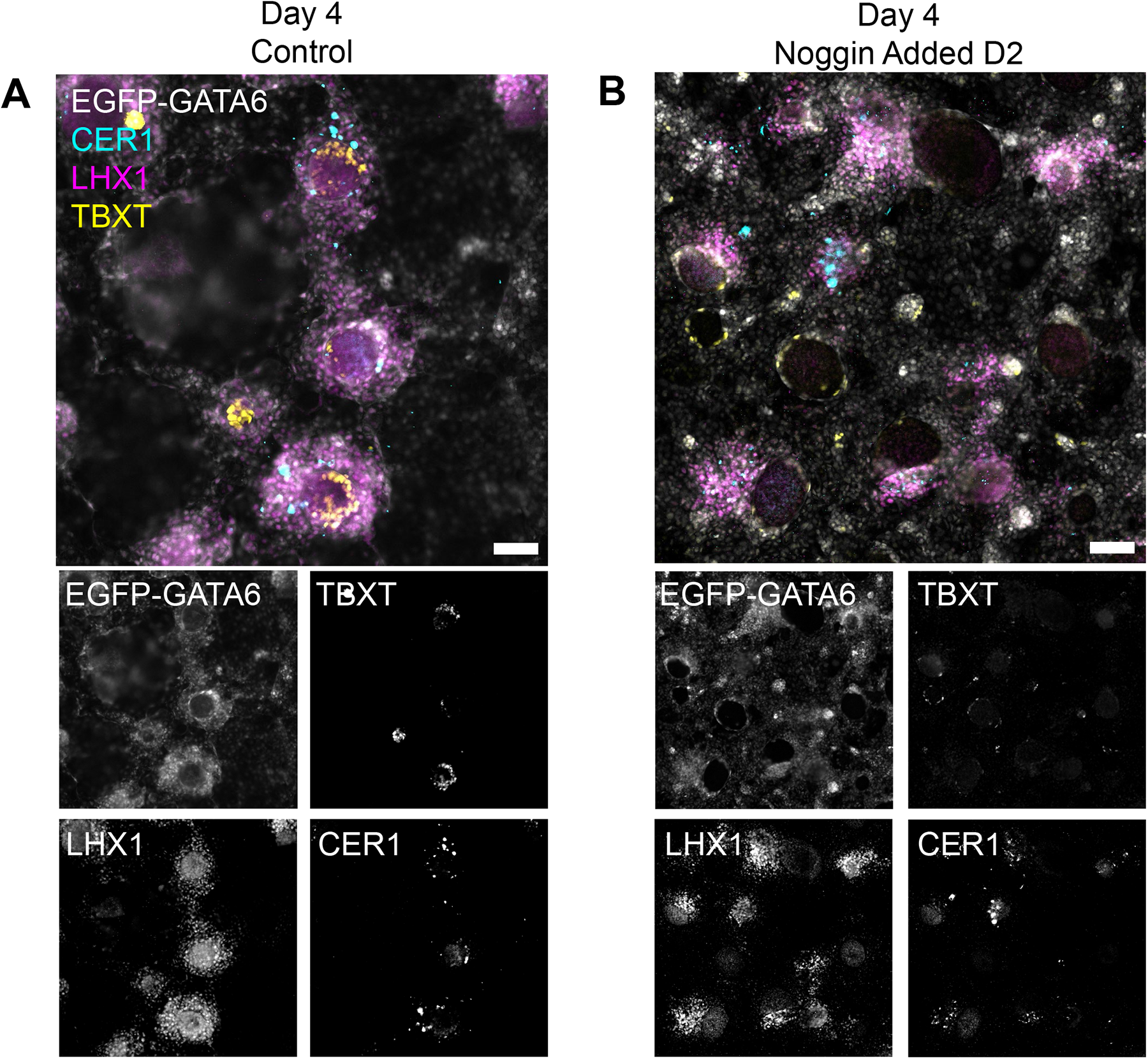
Effect of BMP4 inhibition on anterior-like and posterior domains in iDiscoids. (A) Control iDiscoids showing development of TBXT+ posterior domains and LHX1+ areas expressing CER1. (B) iDiscoids with 100ng/mL Noggin added at day 2 showing presence of LHX1+ areas expressing CER1, but no expression of TBXT+ domains. TBXT expression is seen in iGATA6-lineage cells at the periphery of the WT disc. Scale bars = 100 μm

**Extended Data Figure 10:**
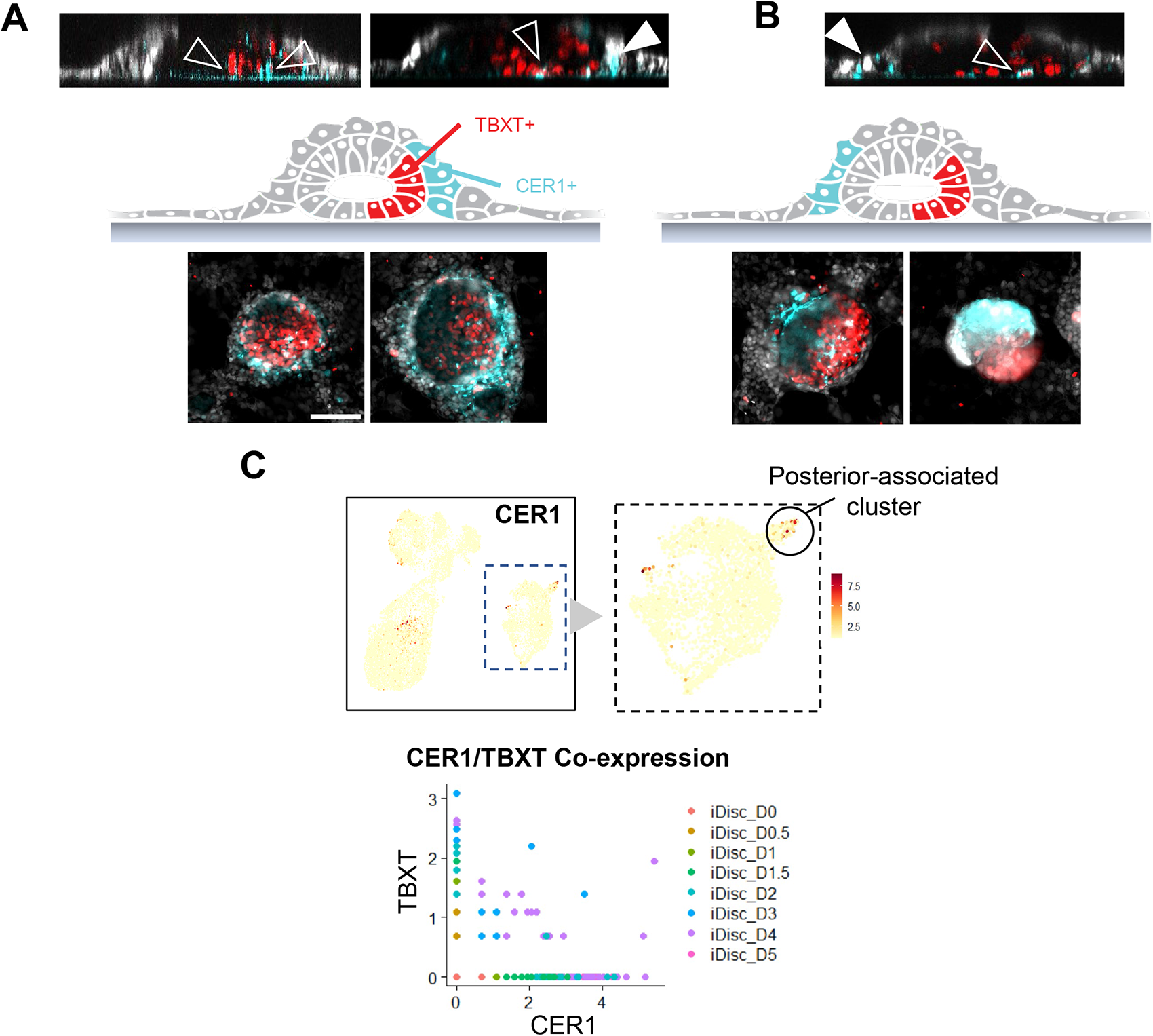
Examples of TBXT and CER1 observed in iDiscoids. (A) A representative WT cluster showing syn-polarity of TBXT and CER1 within the WT cluster and iGATA6 layers, respectively. Filled arrow indicates CER1-expressing cells within the iGATA6 layer; empty arrow indicates CER1 and TBXT co-expressing cells within the WT layer. Z-slices are representative slices from the center of two different WT discs. (B) Representative WT clusters showing anti-polarity of TBXT and CER1 within the WT cluster and iGATA6 layers, respectively. Filled arrow indicates CER1-expressing cells within the iGATA6 layer; empty arrow indicates CER1 and TBXT co-expressing cells within the WT layer. Z-slice is a representative slice from the center of a WT disc. (C) Top shows a merged UMAPs of all D0-D5 scRNA-seq data showing the posterior compartment expressing the inhibitor CER1. Dotted boxes show WT lineages. Posterior region was illustrated in Figure 3B. Bottom shows a scatterplot from merged D0-D5 iDiscoid scRNA-seq showing a population of cells derived from D3-D4 iDiscoid that co-express CER1 and TBXT. Axes represent scaled expression levels of each marker; dots represent individual cells colored by day. Scale Bars = 100 μm

**Extended Data Figure 11:**
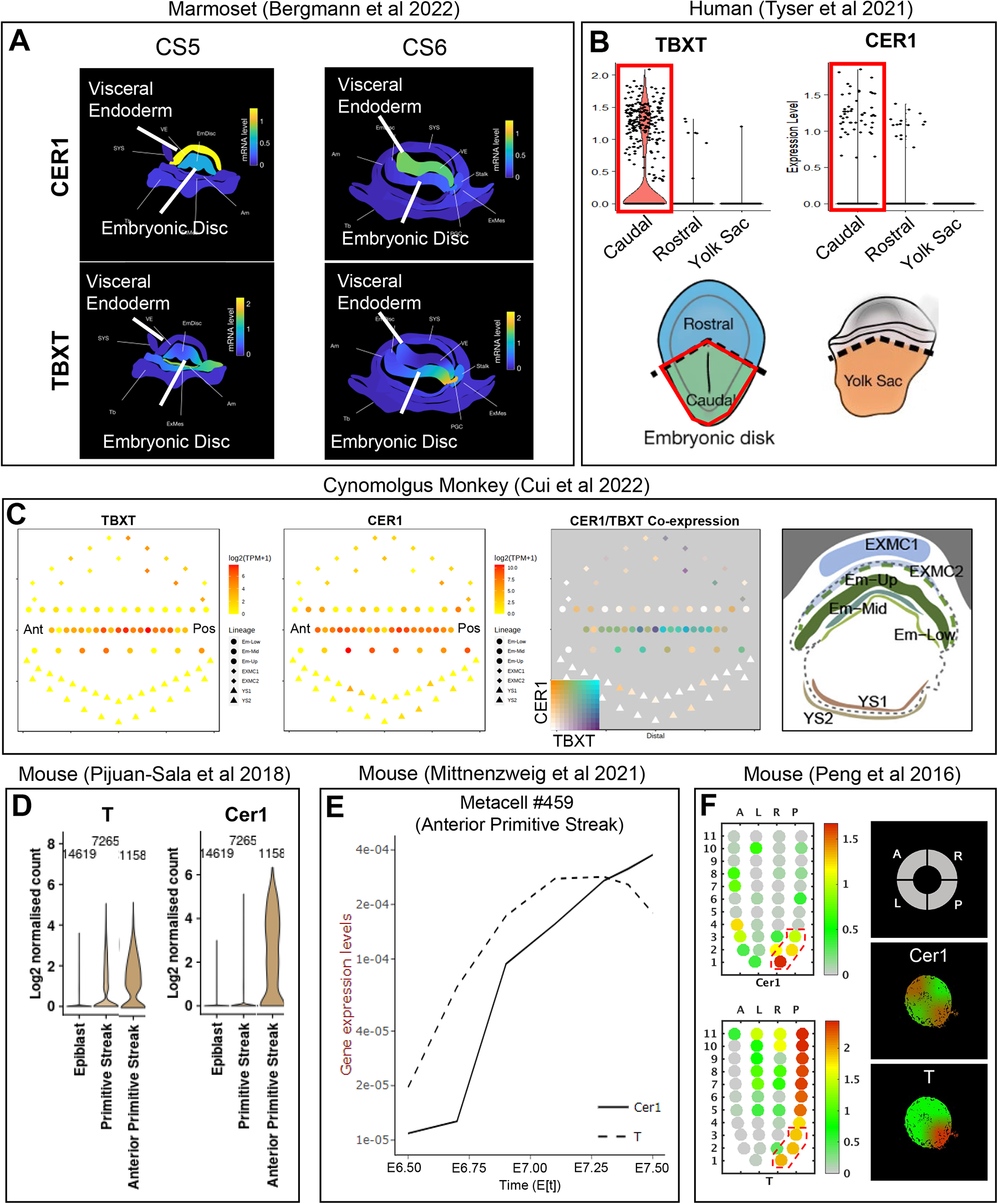
Positioning of TBXT and CER1 expression in transcriptomic human, NHP and mouse datasets. (A) Spatial expression of TBXT and CER1 from the CS5 and CS6 marmoset embryo from Bergmann et al. 2022. In marmoset CS5, CER1 has high and unpolarized expression in the extraembryonic endoderm overlaying the embryonic disc; by CS6, overall expression has decreased, and a very slight polarization away from the putative posterior is observed. From CS5 to CS6, TBXT moves from unregionalized/bipolar expression within the embryonic disc to strong expression at one pole of the embryonic disc. (B) Regional expression of TBXT and CER1 in the E16-19 (CS7) human embryo from Tyser et al. 2022. Highest expression of TBXT and CER1 is observable in the caudal portion of the embryo, which corresponds to the area of the primitive streak and the hypoblast overlaying this area, as shown in the diagram. (C) Pseudospatial representation of TBXT and CER1 expression in the cynomolgus E18-E20 embryo from Cui et al. 2022. Diagram to the right shows the microdissected embryo sections with labels corresponding to each identified tissue layer. CER1 expression strongly overlays TBXT expression in the lower embryonic (EM-Low) and middle embryonic (EM-Mid) regional layers. (D) Expression levels of T and Cer1 from the E6.5-E8.5 mouse embryo. There is a pronounced co-expression within the anterior domain of the primitive streak. (E) Expression levels of T and Cer1 from an anterior primitive streak metacell from 153 mouse embryos spanning E6.5 – E7.5. Within this representative cell/cell type, T and Cer1 rise together over time at the anterior primitive streak. (F) Spatial expression levels of T and Cer1 within the ∼E6.5-E7.5 mouse embryo. These markers are co-expressed in the area near the anterior tip of the posterior domain. Boxed area in the diagram corresponds highlights this range. Right diagrams show expression patterns of each marker at the index 3 plane in the Y axis of left diagrams.

**Extended Data Figure 12:**
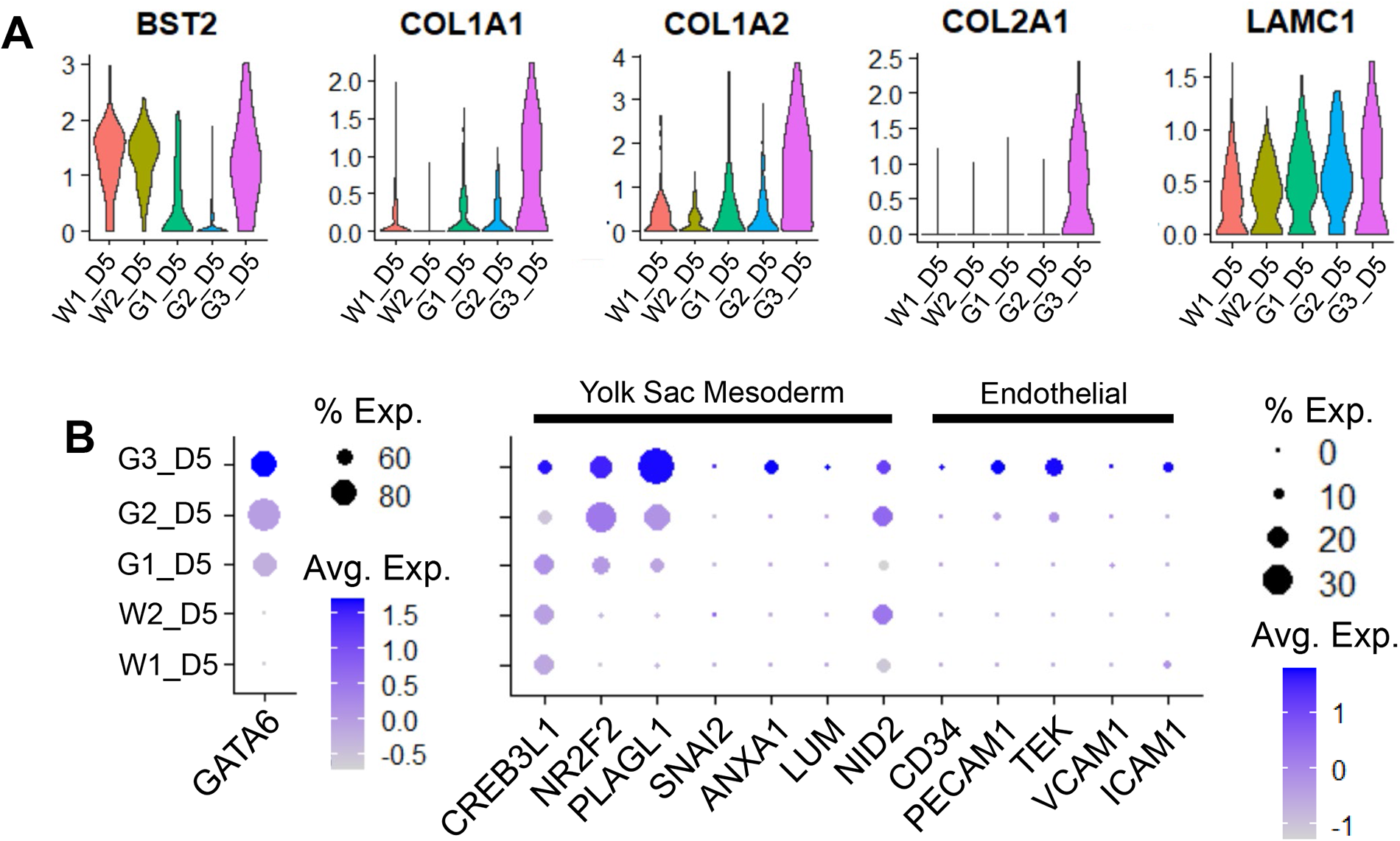
Expression of yolk sac mesoderm and extracellular matrix genes in day 4 iDiscoid scRNA-seq. (A) Violin plots showing the distribution of ECM genes as well as the yolk sac mesoderm marker gene BST2 in day 5 iDiscoid scRNA-seq clusters. (B) Dot plot showing a subset of day 5 yolk sac mesoderm and endothelial marker genes. The highest GATA6 population expresses the highest levels of YSM marker genes. The GATA6 expression is shown to the left with different scaling of circle sizes than the marker list to the right.

**Extended Data Figure 13:**
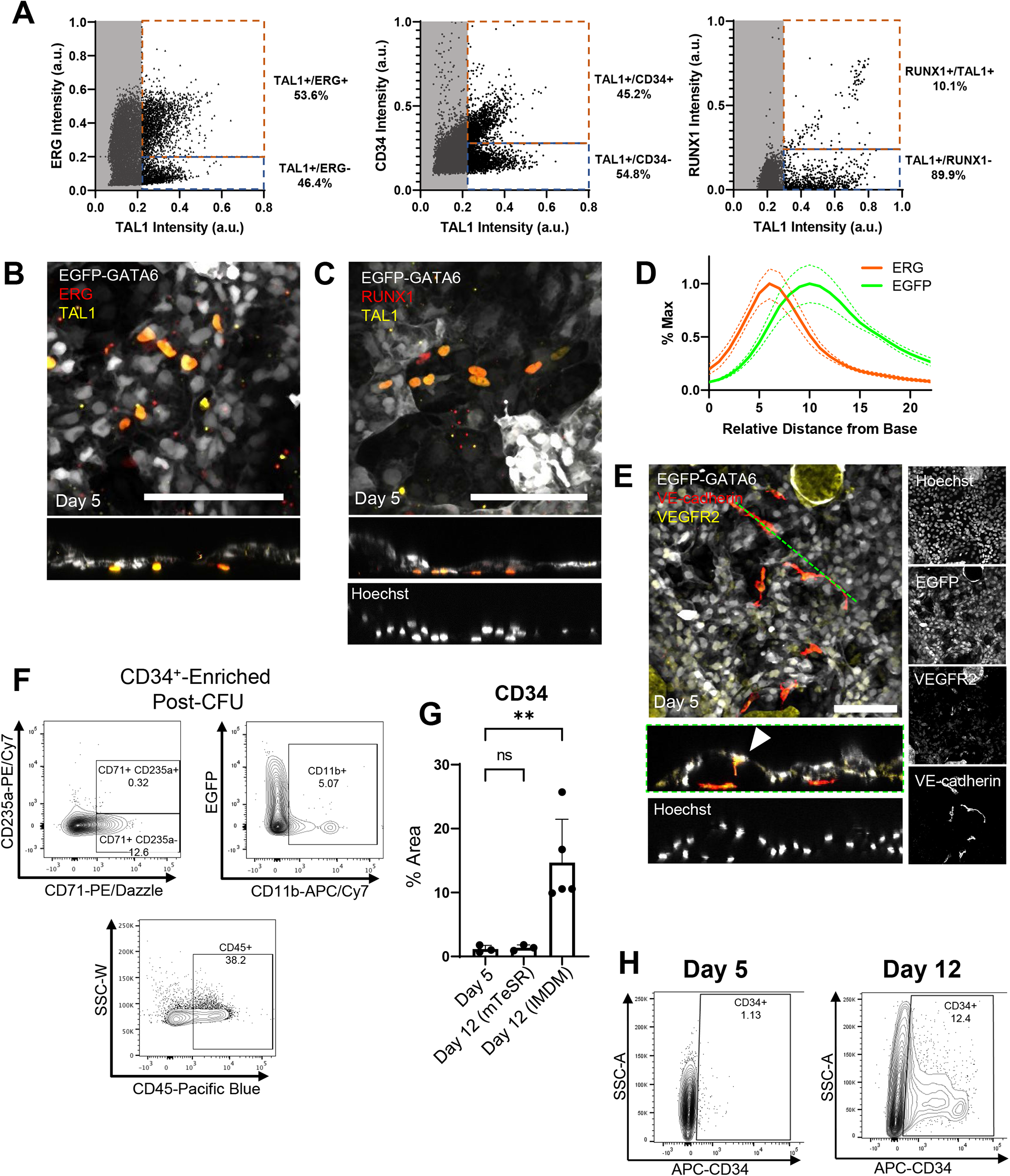
Identification of endothelial/hematopoietic populations. (A) Scatterplots showing the distribution of markers obtained via image analysis of 3 independent experiments at day 5. Percentages correspond to the fraction of cells recorded with corresponding marker expression levels above the thresholds represented by the dotted lines. (B) Immunofluorescence image showing cells expressing hematopoietic markers in iDiscoid culture. Cells expressing the hematopoietic marker TAL1 (Scl) localize between the yolk sac endoderm compartment and the tissue culture dish. Scale bar = 100 µm (C) Immunofluorescence image showing cells expressing RUNX1 and TAL1 positioned against the dish. Scale bar = 100 µm (D) Image analysis of z-slices from 5 areas in 3 biological replicates. The peak of ERG expression is underneath the peak of EGFP expression representing the iGATA6 endoderm layer. Dotted curves represent SEM calculated at each point. (E) Immunofluorescence image showing cells expressing VE-cadherin and VEGFR2 positioned against the dish. Arrow indicates likely differentiating endothelial cell from the overlaying iGATA6 layer. Scale bar = 100 µm (F) Flow cytometry plots showing the presence of hematopoietic markers on day 14 after a CFU assay initiated CD34-expressing cells (enriched from iDiscoids). (G) Results from image analysis demonstrating the change in area of CD34+ cells between day 5 and day 12 of iDiscoid culture, assessed via analysis of immunofluorescence images. Individual dots represent biological replicates. **: P value=0.0055 (H) Representative flow cytometry plot on day 5 and day 12 of iDiscoid showing expansion of the CD34+ population by day 12 (n=3).

**Extended Data Figure 14:**
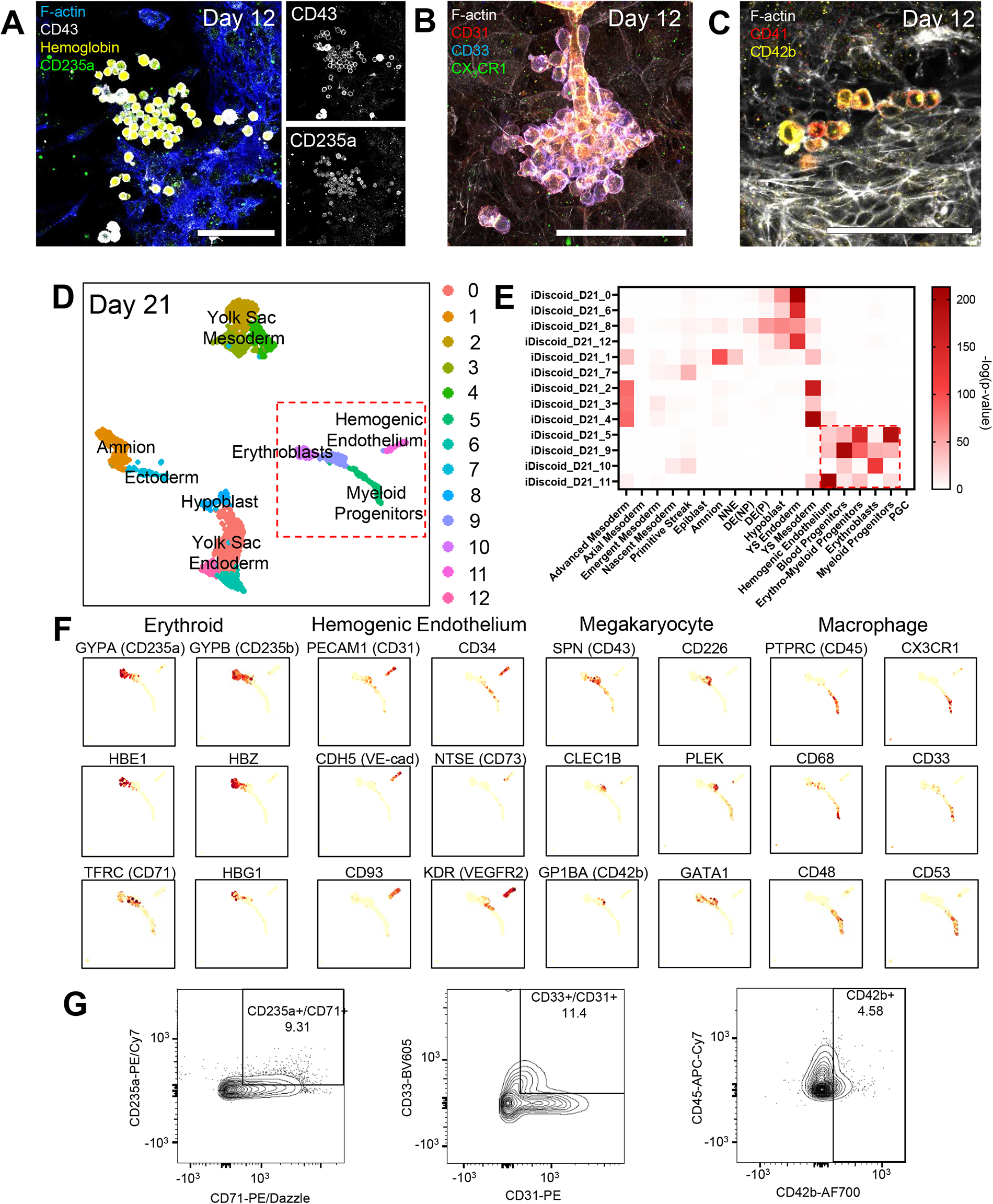
Hematopoietic cells in iDiscoid. (A) Erythroid markers (CD235a and Hemoglobin) expressed in a subset of CD43+ round cells in day 12 iDiscoids. Scale bar = 100μm (B) Cells expressing myeloid (CD31, CD33) and macrophage (CX_3_CR1) markers in a representative cluster in day 12 iDiscoid. Scale bar = 100μm (C) Cells expressing megakaryocyte markers in day 12 iDiscoid. Scale bar = 100μm (D) UMAP showing unbiased clustering of scRNA-seq performed on day 21. Boxed area shows populations expressing hematopoietic marker genes. (E) Hypergeometric statistical comparison of day 21 iDiscoid populations to the E16-19 human embryo populations from Tyser et al. 2022. The red box shows populations of interest with similarity to in vivo hematopoietic lineages. iDiscoid clusters are grouped by similar fates on Y axis. (F) Expression of hematopoietic marker genes within the red boxed population in panel D. The four clusters show distinct marker gene expression profiles associated with yolk sac-derived hematopoietic populations. (G) Flow cytometry analysis on day 21 iDiscoid. Cells were gated for positive CD43 expression prior to gating for the markers shown (n=2, representative sample shown).

**Extended Data Figure 15:**
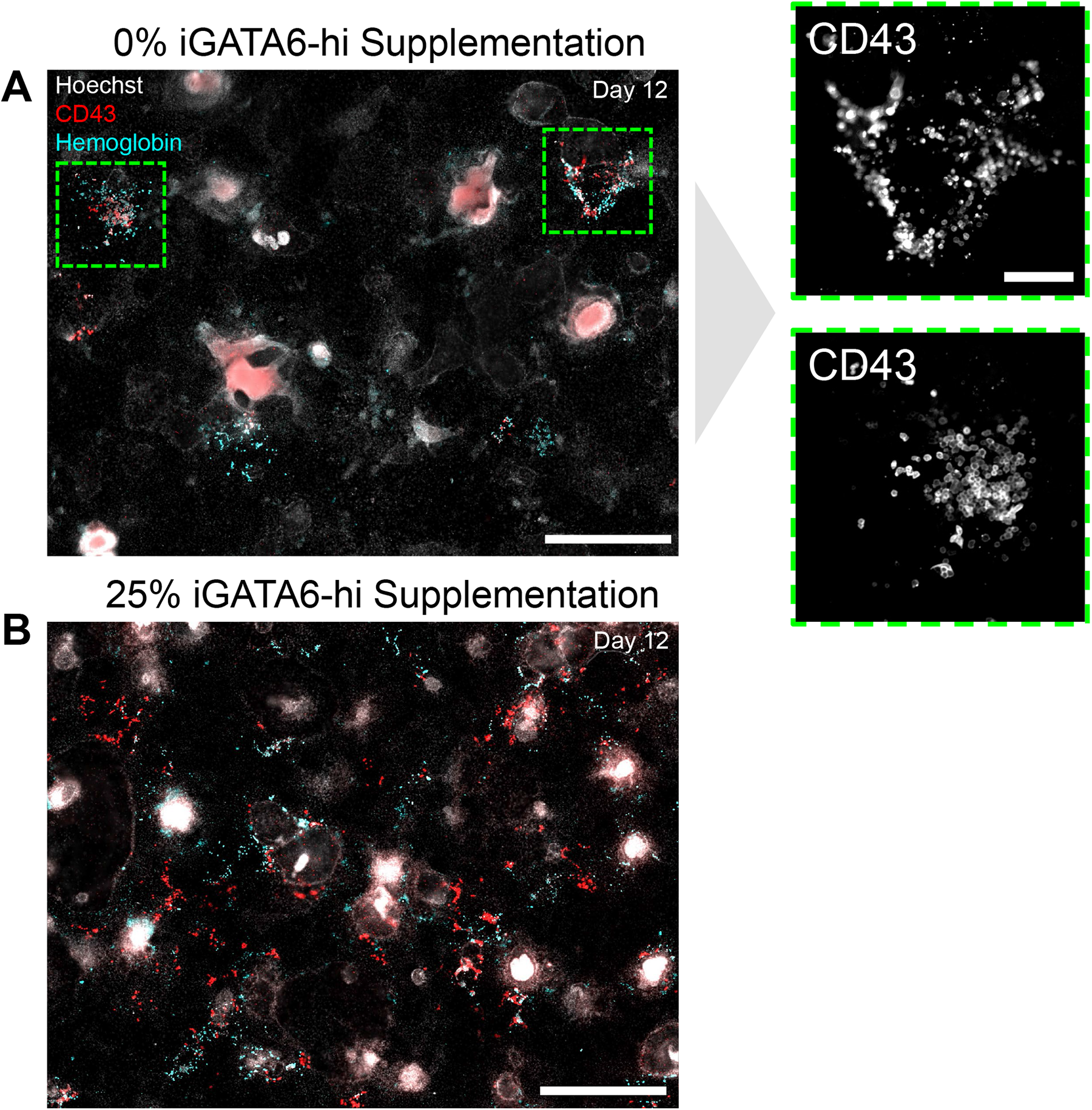
Comparison of iDiscoids with and without supplementation with high iGATA6 expressing (iGATA6-hi) cells. (A) Expression of CD43+ and Hemoglobin+ cells in day 12 iDiscoid without additional supplementation of GATA6-hi cells. Red color in high Hoechst areas is nonspecific staining. Scale bar = 500 μm. Insets show areas of CD43+ cells. Dotted outlines on right show zoomed in areas shown within the image. Scale bar = 100 μm (B) Expression of CD43+ and Hemoglobin+ cells in day 12 iDiscoid with supplementation of 25% GATA6-hi cells at initial seeding. A significant expansion of CD43-expressing cells is observable. Scale bar = 500 μm. A representative image of 2-4 biological replicates.

**Extended Data Figure 16:**
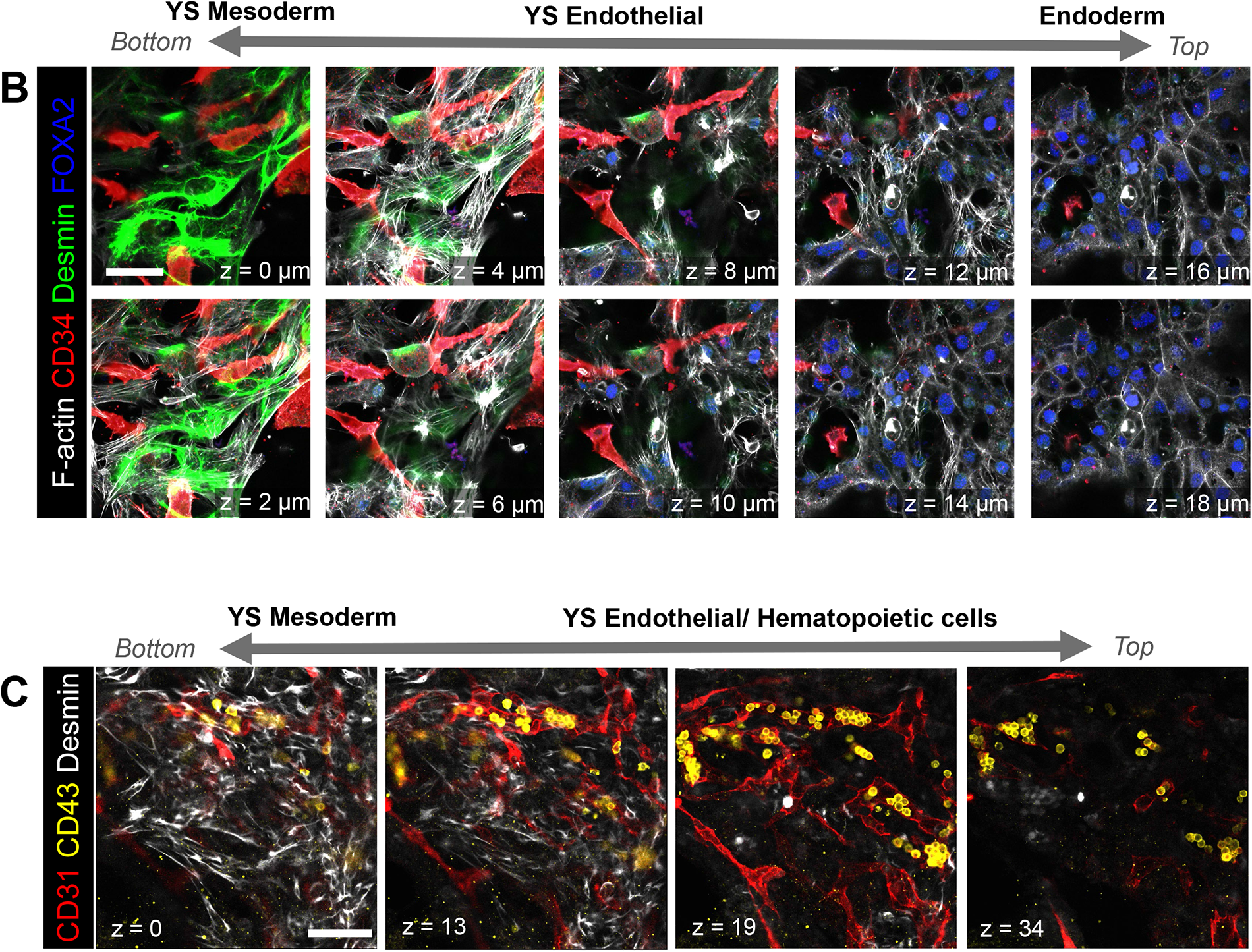
Structure of blood island-like associated tissues within iDiscoid. (A) Z-slices of a day 12 iDiscoid at the indicated height above the bottom focal plane. Expression of the mesodermal marker Desmin is regionalized close to the bottom of the tissue; endothelial marker CD34 is expressed near the bottom and middle of the tissue, with highest expression just above Desmin (z = 6-8); endoderm marker FOXA2 is exclusively expressed above the other tissues. Scale bar = 50 μm (B) Z-slices around the blood island-like area shown in Figure 4E. Desmin is observed regionalized below the endothelial (CD31+) and hematopoietic (CD43+) tissues. No stain for endoderm is shown. Scale bar = 100 μm

**Extended Data Figure 17:**
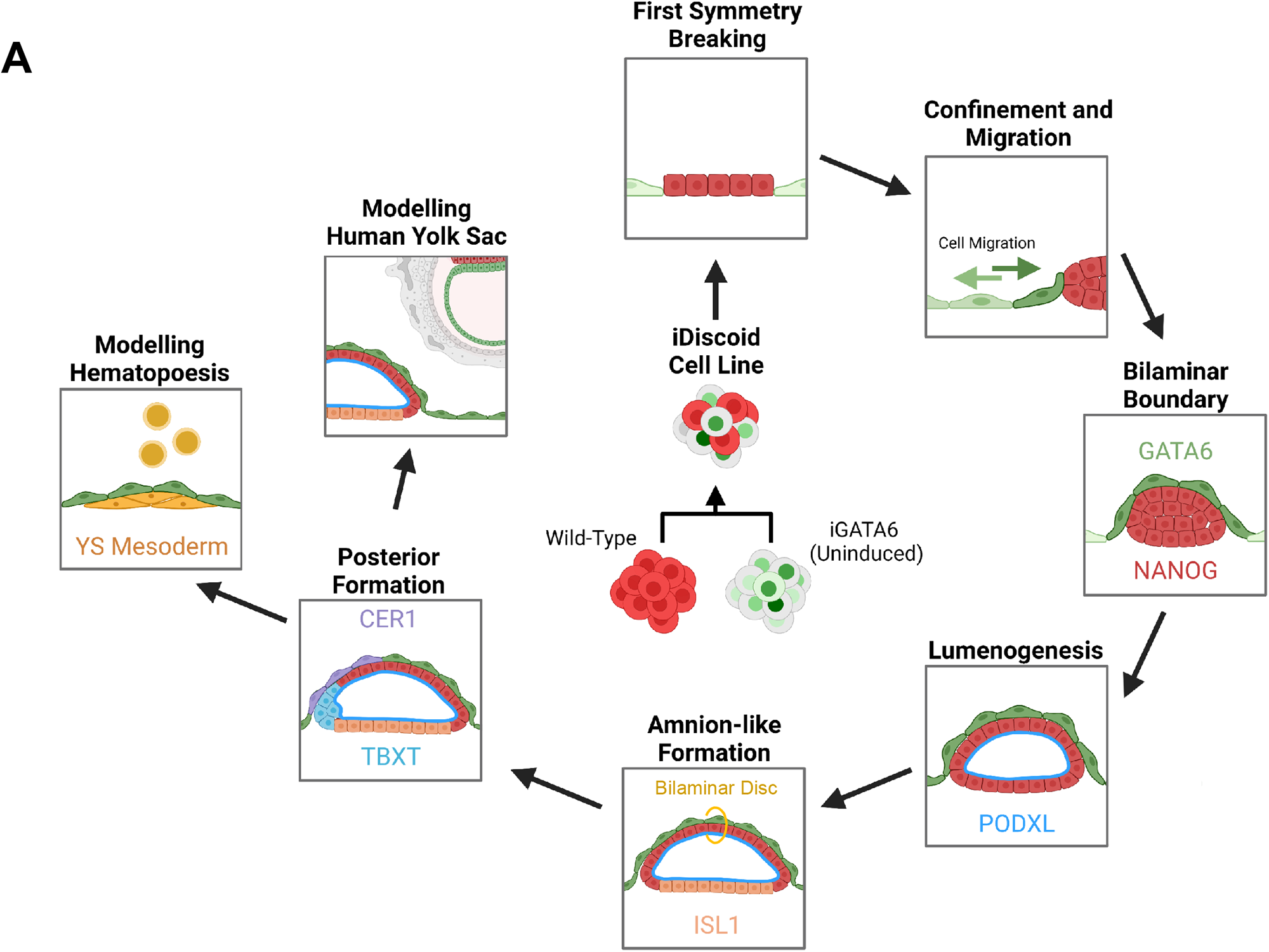
iDiscoid formation from hiPSCs to model human early post-implantation development *in vitro*. (A) From an initially mixed state, iDiscoid cells segregate into WT clusters surrounded by iGATA6 cells. These iGATA6 cells migrate laterally to create a bilaminar boundary on top of the WT clusters. These clusters then undergo lumenogenesis, specification of amnion-like cells, and formation of anterior hypoblast-like and posterior domains. In iGATA6-only areas, yolk sac mesoderm specifies and hematopoietic cell specification is observed.

**Extended Data Figure 18:**
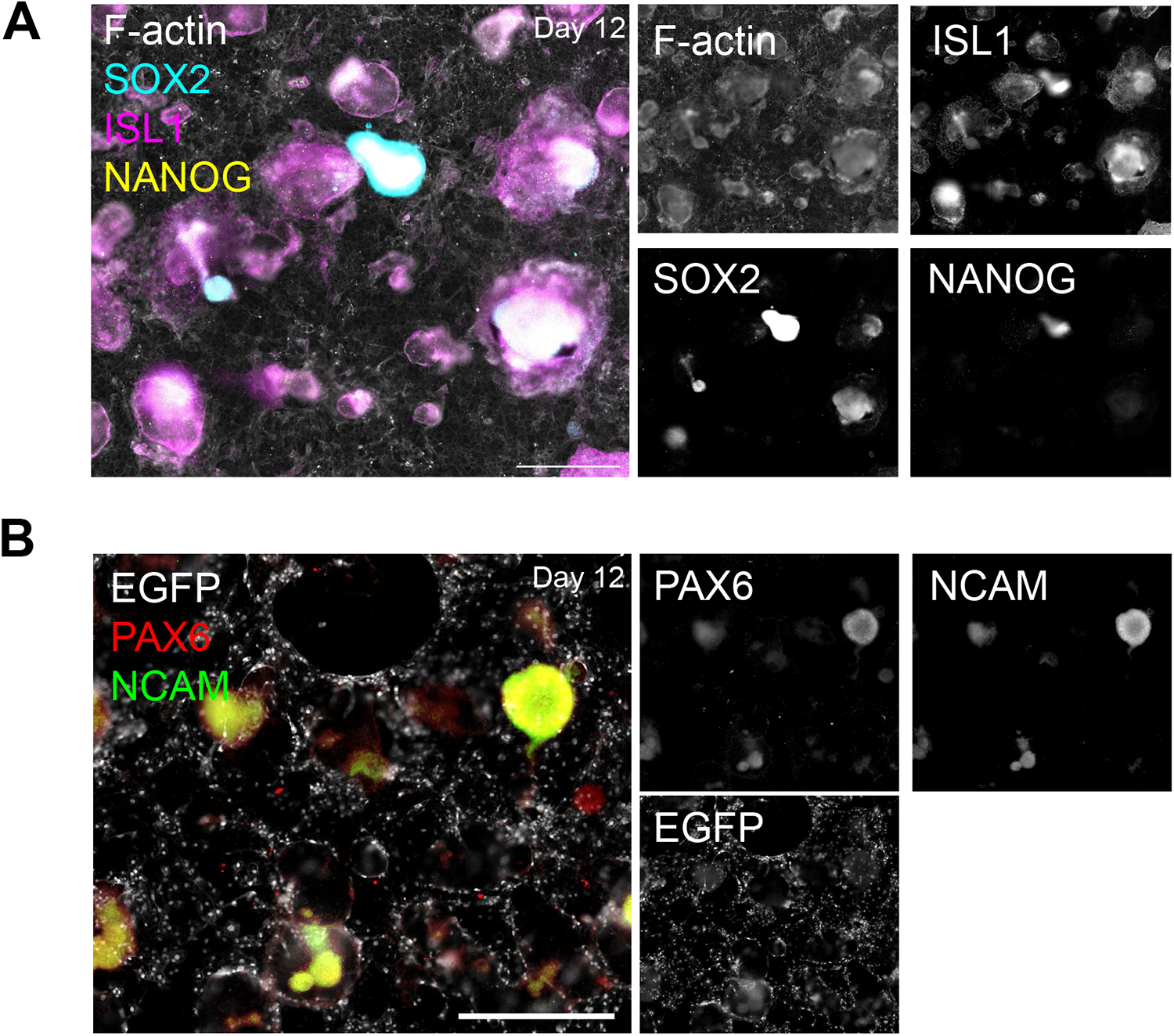
Investigation of WT fates in day 12 iDiscoid. (A) Most structures corresponding to former WT clusters at day 12 in iDiscoid have taken on expression of ISL1 and lost expression of the pluripotency markers SOX2 and NANOG. A limited number of cells express SOX2 at the core of former WT clusters, potentially indicating the specification of an ectoderm-like fate in a small number of WT-lineage cells. (B) A subset of former WT clusters have taken on markers of ectoderm differentiation. Scale bars = 500 μm

**Extended Data Figure 19:**
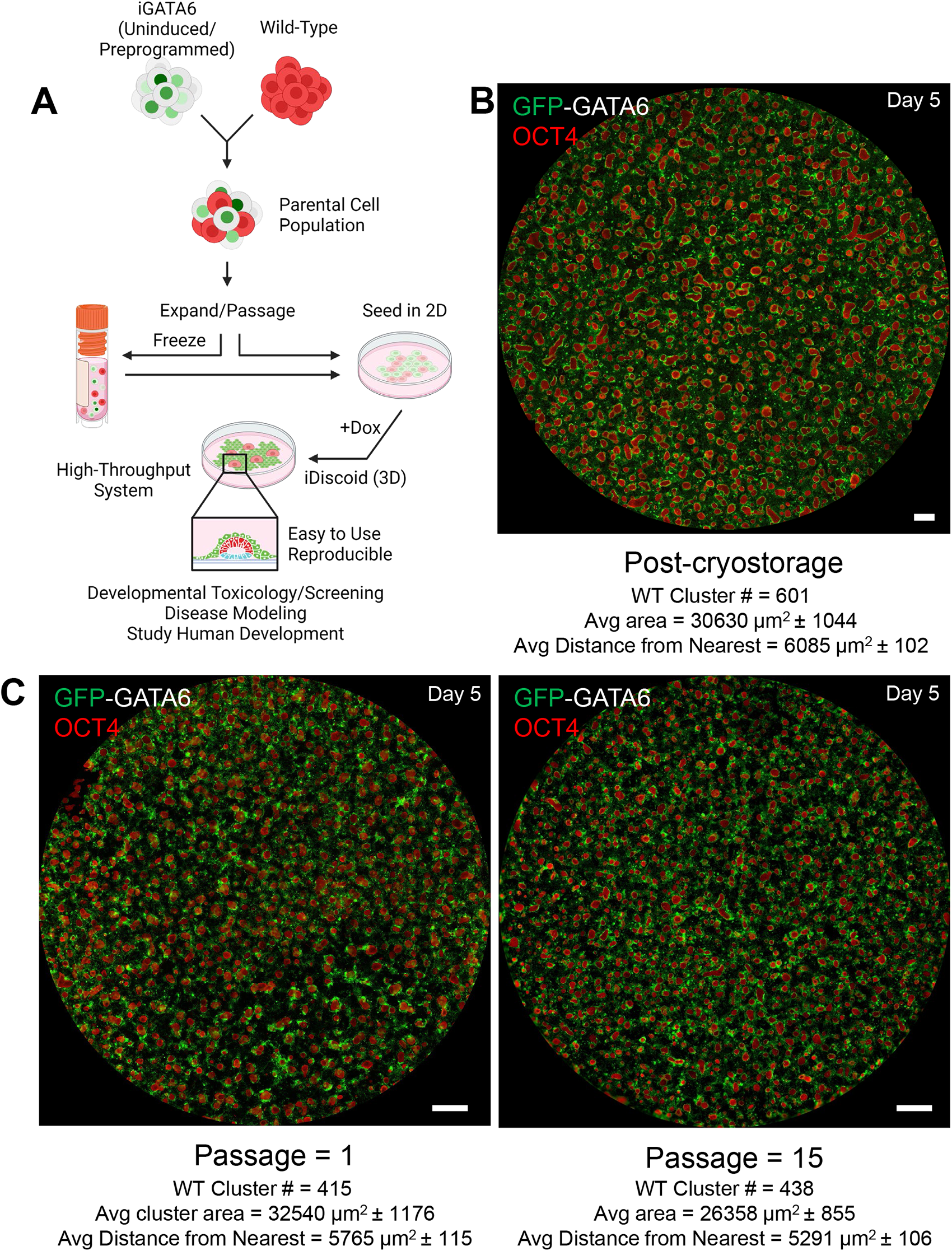
Mixed iDiscoid cell lines exhibit consistent cluster formation after repeated passaging and cryostorage prior to induction. (A) Schematic showing the creation of the iDiscoid parental cell line. iGATA6 cells with heterogeneous copy numbers of the inducible GATA6 circuit are mixed with wild-type. This cell combination is then maintained together or frozen prior to induction for iDiscoid experiments. (B) iDiscoid morphology and characteristics following cryostorage and defrosting. Scale Bar = 500 μm (C) Characteristics and morphology of iDiscoid cultures induced immediately after mixing (passage 1) or maintained together and passaged for two months (passage 15). Scale Bars = 500 μm

**Extended Data Figure 20:**
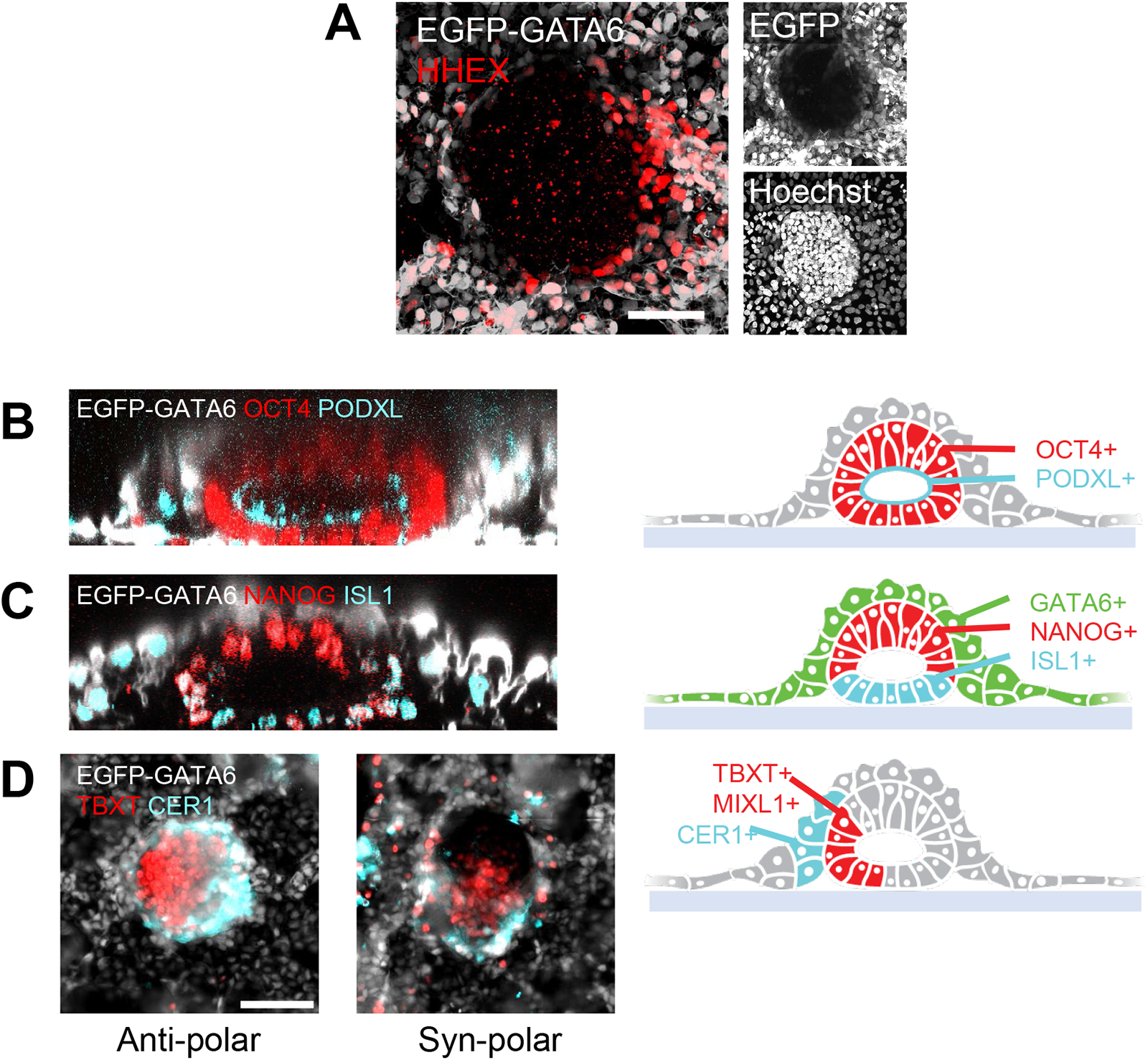
iDiscoid engineering in a separate iPSC cell line results in the generation all major features of interest. Day 4 immunofluorescence images of iDiscoid engineered using the iPSC cell line PGP9. (A) A portion of PGP9 iGATA6 cells expressing high levels of GATA6 (EGFP) also express the anterior endoderm marker HHEX near the edge of a WT cluster. (B) A ring of PODXL expression lines the inside of a cavities formed in PGP9 WT clusters. (C) ISL1+ cells specify away from NANOG+ cells, along the base of a cavity formed in PGP9 WT. (D) TBXT+ cells develop polar domains at the edge of WT clusters. Representative islands show that both syn-and anti-polar arrangement of cells are observable. Scale Bars = 100 μm

**Extended Data Figure 21:**
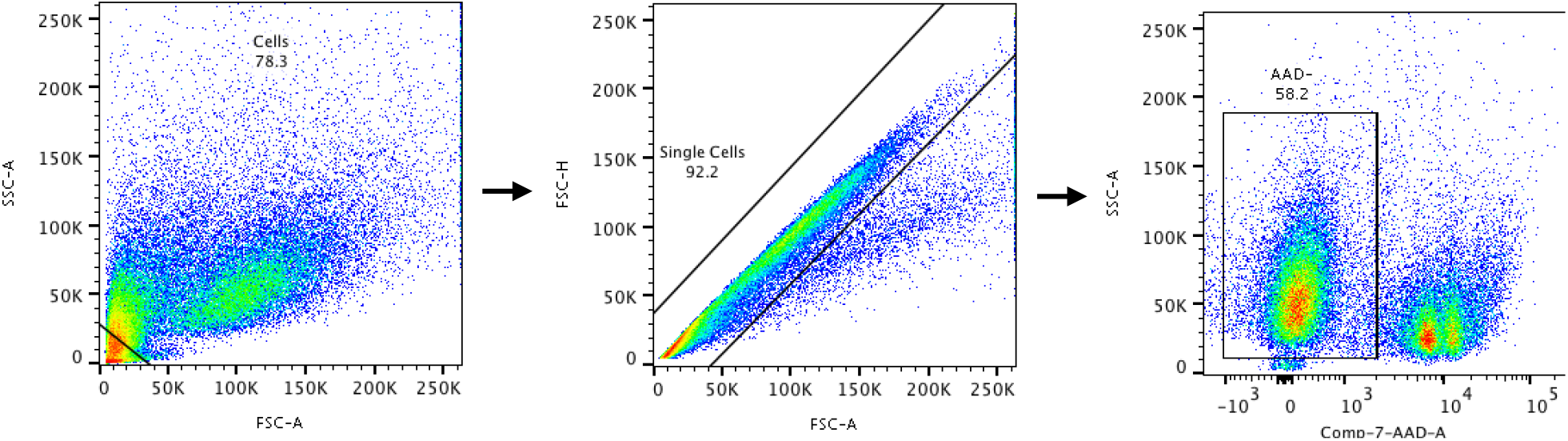
Example of FACS gating strategy. (Cited in Methods) Cell debris was excluded via an SSC-A vs FSC-A gate; aggregates were excluded by comparing FSC-A and FSC-H; dead cells were gated out using the 7-AAD stain to identify positive cells.

## Supplementary Information

**Supplementary Video 1: Self-organization of hiPSC cells that result in WT discs surrounded by iGATA6 cells from day 0 to day 3**

**Supplementary Video 2: Migration of iGATA6 cells over the top of the WT cells to form a contiguous membrane** iGATA6 cells (green) cover the top of the WT disc (red) between day 0.5 and day 2.

**Supplementary Video 3: Expansion of a lumen from a rosette in a single iDiscoid**.

**Supplementary Video 4: Fly-through of an iDiscoid with a single lumen** Z-slices of a single fixed iDiscoid moved through from end-to-end. DAPI: blue, iGATA6: Green, PODXL: Red, F-actin: Gray

